# BET BD2 inhibition facilitates SPOP-mediated degradation of chromatin-associated BRD4/BRD4-NUT, a therapeutic vulnerability in NUT carcinoma

**DOI:** 10.64898/2026.08.14.744905

**Authors:** Kathleen A. Bates, Hoang Nguyen, Kyle P. Eagen, Julianna Huang, Prafulla C. Gokhale, Brittaney A. Leeper, Benjamin K. Eschle, Stacey T. Gray, Krupa Sampat, R. Taylor Durall, Jia Luo, Geoffrey I. Shapiro, Steven J. Ferrara, Jacob H. Gillis, Dustin Rogers, Kfir R. Schreiber, Luca Rastelli, Madeleine E. Lemieux, Christopher A. French

## Abstract

BET bromodomain inhibitors block binding of BET family bromodomains 1 and 2 (BD1, BD2) to chromatin and have demonstrated clinical activity in NUT carcinoma (NC), a BRD-NUT fusion-driven cancer, but toxicity from BD1 inhibition has limited their effectiveness. We investigated whether selective inhibition of BRD4 bromodomain 2 (BD2) could retain antitumor activity while reducing toxicity. NC cells were uniquely sensitive to the novel BRD4-BD2 inhibitor DC-9476 and other BD2-selective inhibitors, which induced differentiation and growth arrest. A CRISPR knockout screen identified the BRD4-targeting E3 ligase SPOP as the top resistance hit. BD2 inhibition, but not BD1-selective or pan-BET inhibition, triggered SPOP-dependent proteasomal degradation of BRD4 and BRD4-NUT; SPOP loss prevented degradation and largely rescued BD2 inhibitor-induced differentiation and growth arrest. Unexpectedly, BRD4 and BRD4-NUT remained chromatin-associated during BD2 inhibition, whereas BD1 or pan-BET inhibition displaced them. Together with evidence that ectopic BRD4-NUT expression sensitizes BRD4 to degradation, these findings support a model in which BRD4-NUT megadomains create a high-density, degradation-competent SPOP substrate pool of BRD4 and BRD4-NUT upon BD2 inhibition, whereas pan-BET inhibition disperses this substrate and limits efficient degradation. In preclinical NC models, BD2-selective inhibition achieved greater tumor growth inhibition and survival benefit than pan-BET inhibition, revealing a therapeutic vulnerability.

## Introduction

NUT carcinoma (NC) is an aggressive fusion oncoprotein-driven squamous carcinoma of the lungs and head and neck^1^. With a median overall survival of 6.7 months, it is one of the most aggressive solid tumors in humans, rivaling metastatic pancreatic cancer^1–4^. Most patients are adolescents and young adults (AYA), and there exist no reliably effective therapies, thus this AYA cancer, though rare, is an extreme unmet need.

NC is driven by NUT-fusion oncoproteins encoded by a variety of chromosomal-translocation-fused genes to *NUTM1*, most commonly (∼75%) *BRD4::NUTM1*^5,6^. Other major fusions include *BRD3::NUTM1* (10-15%)^2,6,7^ and *NSD3::NUTM1* (10-15%)^2,6,8^, followed by exceptionally rare variants fusing ZNF-encoding genes to *NUTM1*^2,3,6,9,10^. All NUT fusion partners are enriched within the NC oncogenic protein complex and bind to the extraterminal (ET) domain of BRD4, thus NUT-fusions have a unifying mechanism to tether NUT to BRD4^9,11^. Because they function similarly, in this report we collectively term BRD3- and BRD4-NUT fusions “BRD-NUT oncoproteins”.

NC is an epigenetic cancer driven entirely by BRD4-NUT ^12,13^, or other BRD-NUT oncoproteins, with few if any additional oncogenic mutations^14,15^. It is characterized by massive BRD-NUT-, p300-, and acetyl-histone-enriched super-enhancers, called megadomains, that upregulate oncogenic genes, namely *MYC* ^9,16,17^. The maintenance of MYC expression in NC ensures continued growth and arrested differentiation^7,16,17^. An important downstream target of MYC is *EZH2*, encoding a key component of polycomb repressor complex 2 (PRC2) which represses the classic tumor suppressor gene, *CDKN2A*^18^, thus bypassing the need for genetic deletion of this gene.

BRD4-NUT is a chromatin reader-writer: BRD4, a BET family protein, binds (“reads”) acetyl-histones with its dual bromodomains. NUT recruits and potently activates the p300 histone acetyltransferase to further acetylate (“write”) local histones ^16,19^. This feed forward process of iterative reading and writing by BRD4-NUT forms the massive megadomains that characterize this disease^16^.

First generation BET bromodomain inhibitors (BETi) are acetyl-lysine mimetics that competitively inhibit both BET bromodomains, termed BD1 and BD2, including those of BRD2-4, and -T, from binding acetyl-histone post-translational modifications (PTM). Here we term these pan-BETi, ‘BD1-BD2-BETi’. We and colleagues have shown that BD1-BD2-BETi evict BRD-NUT from chromatin, resulting in downregulation of MYC, induction of squamous differentiation and arrested growth^20^. We and others have also shown that BD1-BD2-BETi can be used therapeutically in NC patients, demonstrating on-target activity with objective response rates from phase 1 trials of 20-30%. Clinical activity of first generation BD1-BD2-BETi has been modest, limited by hematologic (thrombocytopenia) and gastrointestinal toxicity^21–24^.

The limited tolerability to BD1-BD2-BETi is believed due to inhibition of BD1, which is required for promoter activation of essential housekeeping genes, which are regulated at the promoter level rather than by enhancers ^25–27^. BD2, in contrast, along with BD1 is involved with enhancer activation of inducible genes, such as *MYC* and those involved in inflammation, and is generally not involved with maintenance of transcription of steady state genes ^26–28^. Thus, to mitigate toxicity, second generation BD2-selective inhibitors (BD2i) were developed, and indeed this class of compounds have demonstrated much less toxicity in animals than BD1-BD2-BETi. However, anti-neoplastic activity of BD2i was limited to only a small subset of cancer types^29,30^.

In this study, we show that BD2i, including the novel BRD4-BD2-selective BETi DC-9476, potently inhibit NC cells regardless of fusion partner, phenocopying the effects of BD1-BD2-BETi. We find that the exquisite sensitivity of NC to BD2i is largely mediated by SPOP-dependent proteasomal degradation of wild type BRD4 and BRD4-NUT. These findings provide a mechanistic rationale for the therapeutic development of BD2i in NC.

## Methods

### Tumor cell lines

NC cell lines, TC-797^31^, 10–15^17^, 14169^32^, and non-NC cell lines, 293T, U2OS (RRID: CVCL_0042), H520 (RRID:CVCL_1566), HCC95 (RRID:CVCL_5137), and Calu-1 (RRID:CVCL_0608) were maintained as monolayer or suspension (10326^7^) cultures in DMEM (Invitrogen) supplemented with 10% FBS (Sigma-Aldrich batch number 24K006, St. Louis, MO, RRID:SCR_008988), 1X GlutaMAX (Thermo Fisher Scientific, RRID:SCR_008452) and 1% pen– strep (Hyclone). PER-403 cells were cultured as monolayers in DMEM (Invitrogen) supplemented with 20% BGS (Hyclone batch number AL30815049, South Logan, Utah) and 1% pen–strep (Hyclone). NCI-H596 [H596] (ATCC HTB-178) cells were cultured as monolayers in RPMI-1640 Medium (ATCC 30-2001) supplemented with 10% FBS (above) and 1% pen-strep (Hyclone). The TC-797, PER-403, 14169 and 10–15 cell lines were authenticated by fluorescent *in situ* hybridization as described^8^ demonstrating rearrangement of the *NUTM1* and *BRD4* genes, and by STR profiling. 10326 cell line was authenticated by fluorescent *in situ* hybridization as described^8^ demonstrating rearrangement of the *NUTM1* and *BRD3* genes, and STR profiling. The 293T, U2OS, H520, HCC95, and Calu-1 cell lines have been authenticated by STR profiling.

### Immunohistochemistry

Formalin-fixed, paraffin-embedded tumor samples from mice were prepared using standard methods^7^. All immunohistochemistry (IHC) was performed on the Leica Bond III automated staining platform. Antibodies and conditions used are listed in **Supplementary Table S1**.

### Quantification of NUT immunohistochemical staining

NUT immunohistochemical staining was quantified using QuPath v0.5.1 (RRID:SCR_018257) on whole-slide brightfield H-DAB images. Viable tumor regions were manually annotated, excluding necrosis, artifacts, and non-tumor tissue. Stain vectors were estimated and held constant across slides. Nuclei were detected using the hematoxylin channel, and nuclear NUT staining was quantified by DAB optical density using fixed thresholds applied uniformly across all samples. The percentage of NUT-positive tumor nuclei was determined in a separate threshold-based analysis by classifying nuclei as positive or negative for nuclear DAB staining, without stratifying positive cells by staining intensity. In parallel, an H-score was calculated by classifying nuclei as negative, 1+, 2+, or 3+ based on nuclear DAB staining intensity, using the formula [1 × (% 1+ nuclei)] + [2 × (% 2+ nuclei)] + [3 × (% 3+ nuclei)], producing a score from 0 to 300.

### Hemacolor cytologic staining

Adherent cells were grown on 25mm glass coverslips (Electron Microscopy Sciences, Morgantown, PA) in 6-well format and stained using the Camco Stain Pak according to the manufacturer’s instructions (Cambridge Diagnostic Products, Inc. , Fort Lauderdale, FL).

### SA-β-gal cytologic staining

Adherent cells were grown on 25mm glass coverslips (Electron Microscopy Sciences, Morgantown, PA) in 6-well format and treated with 1µM DC-9476 for 144h. Post treatment, coverslips were fixed and stained overnight using Senescence beta-Galactosidase Staining Kit (CST #9860). The stained coverslips were then counterstained with 125µL Hematoxylin (Fisher healthcare #220-100).

### Quantification of senescence-associated β-galactosidase staining

SA-β-gal staining was quantified from bright-field images using Fiji/ImageJ. β-galactosidase-positive cells were identified by color thresholding in Lab color space using fixed pass ranges of 0–192, 0–136, and 0–109 for the three Lab channels, respectively. Thresholded objects with an area ≥6 pixels² were counted using the Analyze Particles function. The same β-gal color-thresholding parameters were applied to all images and conditions.

Total cell numbers were independently quantified from the hematoxylin counterstain. Images were split into RGB channels and the green channel was used for cell segmentation. Background was removed using the rolling-ball algorithm with the “light background” option, followed by fixed intensity thresholding, conversion to a binary mask, binary opening and closing, watershed separation, and particle counting. Segmentation parameters were optimized by visual inspection to minimize cell fragmentation, merging, and inclusion of debris and were subsequently held constant for all images within each experimental condition. For 10-15 cells, a rolling-ball radius of 30 pixels was used with thresholds of 0–232 for DMSO-treated cells and 0–246 for DC-9476-treated cells, which are larger and flatter; particles ≥6 pixels² were counted. For 14169 cells, DMSO-treated cells were analyzed using a rolling-ball radius of 30 pixels, a threshold of 0–244, and a minimum particle size of 6 pixels², whereas DC-9476-treated cells were analyzed using a rolling-ball radius of 50 pixels, a threshold of 0–246, and a minimum particle size of 50 pixels² because of the larger cell size and increased debris in this condition. These condition-specific parameters were used solely to determine the total-cell denominator and did not influence classification of cells as SA-β-gal positive.

For each biological replicate, the percentage of SA-β-gal-positive cells was calculated as the total number of β-gal-positive cells divided by the total number of cells analyzed ×100. Because SA-β-gal-positive cells were rare in 10-15 cells, five fields were pooled within each biological replicate, yielding total cell counts of approximately 14,500–45,000 cells per replicate. For 14169 cells, in which SA-β-gal positivity was substantially more frequent, one representative field was analyzed from each biological replicate, yielding approximately 1,500–7,200 cells per replicate. Images and fields were treated as subsamples and were not considered independent replicates. Three independent biological replicates were analyzed per treatment condition. DMSO- and 1 µM DC-9476-treated samples were compared using a two-tailed paired Student’s t-test, with each independent experiment constituting a matched pair. Individual biological replicate values are shown together with the mean ± SD.

### Immunoblotting

For standard immunoblots assaying expression of non-histones and non-histone-associated proteins, cell lystes were prepared as described^33^. Briefly, cells were lysed in RIPA Buffer (50 mM Tris-HCl, 250 mM NaCl, 1% NP-40, 0.5% Sodium Deoxycholate, 0.1% SDS, 5 mM EDTA) containing 250 mM NaCl and Halt Protease and Phosphatase Inhibitor Cocktail (78447, Thermo Fisher Scientific) and PMSF (93482, Sigma Aldrich) for 30 min at 4°C with rotation. The lysate was centrifuged at 16,100 x g, and the supernatant was collected.

For assessing expression of histone and histone-associated proteins, whole cell lysates, including chromatin, were prepared as follows and as described. Cells were lyzed in RIPA buffer (50 mM Tris-HCl, 150 mM NaCl, 1% NP-40, 0.5% Sodium Deoxycholate, 0.1% SDS) supplemented with 1 mM EDTA, PMSF, and 2mM MgCl_2_ for 20 min at 4degC. To inhibit protease activity without the addition of Halt Protease and Phosphatase Inhibitor Cocktail, lyzed cells were then incubated at 95degC for 5 min. To obtain complete digestion of DNA and release of its associated proteins into solution, recombinant Dr. Nuclease (Syd Labs, Inc. Natick, MA) was added 1:100 to each sample and incubated at 37degC for 30 minutes, mixing thoroughly. Immunoblotting for both lysis methods was performed as described previously^34^. Antibodies and conditions used are listed in **Supplementary Tables S2-3.**

### CSK-based chromatin fractionation

Nuclear fractionation of cellular compartments was employed to compare protein expression between soluble and insoluble proteins. Cells were lysed in CSK-150 buffer (10mM HEPES (pH 7.4), 150mM NaCl, 300mM sucrose, 3mM MgCl_2_, 1mM EGTA, 0.5% TritonX-100, and 1mM DTT) supplemented with Halt Protease and Phosphatase Inhibitor Cocktail and PMSF for 5 minutes on ice after thorough mixing. Soluble and insoluble proteins were then separated by differential centrifugation (2000xg for 5 minutes at 4°C). The supernatant was collected as the soluble fraction. The remaining pellet was resuspended in SDS-containing sample buffer and sonicated using a Q125 Sonicator (QSONICA 4422, Newtown, CT; RRID: SCR_019046) to become the insoluble fraction. Four-second-long pulses (25% Amplitude) were performed in duplicate, with 10-second rests between.

### Immunofluorescence

Immunofluorescence was performed on adherent cells grown on 25mm glass coverslips (Electron Microscopy Sciences, Morgantown, PA) in 6-well format. Cells were fixed using 4% paraformaldehyde and nuclei were counterstained with ProLong Gold antifade reagent with 4′,6-diamidino-2-phenylindole (DAPI; Life Technologies). Primary and secondary antibodies are listed in **Supplementary Tables S4-5**. Photographs were taken on a Nikon Eclipse E600 fluorescent microscope using a Spot RTSlider camera (Diagnostic Instruments, Inc.), and Spot Advanced software (Diagnostic Instruments, Inc.). For each antibody, exposure time and all other acquisition settings were held constant across conditions. Post-acquisition image processing, including levels adjustments, was performed identically for all images within a given antibody channel.

### Plasmids, cloning, and viral transduction

Creation of stable lentiviral Cas9-expressing 10-15 cells were described previously^18^.

pRDA-355, a lentiviral sgRNA-guide-only, doxycycline-inducible vector was purchased from Addgene (Plasmid # 187159, Watertown, MA, RRID:Addgene_187159) and used to create NC cell line derivatives to knock out BRD4L and SPOP using two constructs each. Creation of pRDA-355 puro non-targeting negative control constructs (sgnegCTRL-1, GGCTTACGTGGGGGGCAAAA (from Brunello library guide, BRDN0001146436^35^); sgnegCTRL-2, GCGAGGTATTCGGCTCCGCG (lacZ targeting)) and constructs targeting SPOP (sgSPOP-1, ACGGGCTTCTCCCTGATGAC; sgSPOP-2, GTGGCCCCGTAGCTGAGAGT) and BRD4L (sgBRD4L-1, GCTGGGTGAAGTGGCCGATG; sgBRD4L-2, GAGGAGAGACCACTGCGTGC) lentiviral plasmids were created by ligating duplexed oligos encoding sgRNAs (synthesized by Integrated DNA Technologies, Inc. (IDT), Coralville, IA) into XhoI digested pRDA-355 using T4 DNA Ligase according to the manufacturer’s instructions (New England Biolabs, Ipswich, MA).

To create lentivirus, 293T (RRID: CVCL_0063) cells were co-transfected with pRDA-355, psPAX2 (RRID:Addgene_12260), and pMD2.G (RRID:Addgene_12259) plasmids using Lipofectamine 2000 (Invitrogen, Waltham, MA). Packaged virus was collected 48h after transfection and purified using a 45 µm pore filter. Polybrene (8 µg/ml, Sigma) was added to viral supernatant and used to transduce 10-15-Cas9 cells. Infected cells were selected using puromycin (1 µg/ml; Sigma-Aldrich (St. Louis, MO) and blasticidin (7.5ug/mL Gibco, Billings, MT) for at least one week. Single clones were isolated and expanded for all experiments described.

pcDNA5 frt/to-*GFP-BRD4-NUTM1*, -*GFP-BRD4-NUTM1*ΔADTTT, and -*flag-BRD4-NUTM1-HA* tetracycline-inducible plasmids were created by LR gateway cloning pDONR221-*BRD4-NUTM1* constructs into a gateway-compatible derivative of pcDNA5 frt/to (pcDNA5 frt/to vector was a gift from Dr. Jon Aster; pDONR221 was a gift from Dr. Mitzi Kuroda). The pDONR221-*BRD4-NUTM1* constructs were created by Gibson assembly, as were pcDNA5-frt/to destination vectors containing N- and C-terminal GFP, HA, or FLAG tags.

### Generation of stable TRex cell line derivatives

The 293TRex and U2OSTRex cell lines and derivatives were created using Flip-In technology as described previously^17^ and according to the manufacturer’s instructions (Invitrogen) and maintained as above, but with the addition of Hygromycin (150 μg/mL; Sigma-Aldrich, RRID:SCR_008988) and Blasticidin (7.5 μg/mL; Life Technologies, Life Technologies RRID:SCR_008817) to maintain selection of cDNA insert and tet repressor genes, respectively. 293TRex-flag-BRD4-NUTM1-HA, U2OSTRex-GFP-BRD4-NUTM1 and GFP-BRD4-NUTM1ΔADTTT derivatives were generated by recombination with pcDNA5 frt/to-*GFP-BRD4-NUTM1*, -*GFP-BRD4-NUTM1*ΔADTTT, and -*FLAG-BRD4-NUTM1-HA* using Flp-In technology (Invitrogen, RRID:SCR_008452). Cells were selected and maintained with 7.5 μg/mL Blasticidin and 150μg/mL hygromycin, and single-cell clones were obtained for down-stream experiments.

### Cell growth assays (Cell Titer Glo)

Cells were seeded into 96-well plates at a density of 500-1,500 cells per well in a total volume of 90 µl media. Compounds were delivered to triplicate wells using a HP D300e digital dispenser (Hewlett Packard, Spring, TX) at the ICCB-Longwood Screening Facility at Harvard Medical School (Boston, MA). For long term (>96h) treatments, cells were re-fed every four days using an Thermo Multidrop Combi (Thermo Fisher Scientific Inc., Waltham, MA) also at the ICCB-Longwood Screening Facility at Harvard Medical School (Boston, MA). Following a 72-96h or 10-day incubation at 37 °C, cells were lysed, and wells were assessed for total ATP content using a commercial proliferation assay (Cell TiterGlo; Promega Madison, WI). Three biologic replicates were performed. Estimates of IC50 were calculated by logistic regression (GraphPad Prism, Dotmatics, Boston, MA, RRID:SCR_002798).

### Chemicals and compounds

EPZ-6438 (tazemetostat) was generously provided by Epizyme (now Ipsen Bioscience, Cambridge, MA). ABBV-744 and ABBV-075 were kindly provided by AbbVie, Inc. (Chicago, IL, RRID:SCR_010484). DC-9476 was provided by DeepCure Inc.. Dimethyl sulfoxide (DMSO) was purchased from Sigma-Aldrich. Tetracycline was purchased from American Bioanalytical. Doxycycline was purchased from Bio Basics.

### RNA extraction and library preparation for RNAseq

Whole RNA was extracted from live cultured 10-15, 10326, and HCC95 cells treated in biologic triplicate for either 18, 24, 36, or 48hr with the indicated concentrations of vehicle (DMSO) and DC-9476 using the RNeasy Plus kit (Qiagen). 1 µg of total RNA was used for the construction of ribosome-depleted sequencing libraries using the KAPA RNA HyperPrep Kit with RiboErase kit (HMR, Roche) according to the manufacturers’ instructions. ERCC ExFold RNA spike-in controls (Invitrogen; 4456739) were added according to manufacturer’s specifications to 1 µg of purified RNA from each sample to help normalize expression. Ribosomal RNA (rRNA) was depleted using the NEBNext rRNA Depletion Kit (NEB; E6350L) according to manufacturer’s instructions. Libraries were 50bp paired-end sequenced on a NovaSeqX Plus at Admera (Admera Health, South Plainfield, NJ).

### Analysis of RNAseq

Samples used were from 3 cell lines, with all at DMSO (24, 48 hours) or DC9476 (18, 24, 36, or 48 hours): HCC95 cells (n=3 DMSO, n=3 DC9476), 10-15 cells (n=2 or 3 DMSO, n=2 DC9476), 10326 cells (n=3 DMSO, n=3 DC9476). All samples were aligned to the Ensembl human genome (Homo_sapiens.GRCh38.dna_sm.toplevel.fa, RRID:SCR_002344) supplemented with External RNA Control Consortium (ERCC92) spike-in sequences added to normalize the samples. Alignment files were converted to TDF format for visualization with the Integrated Genome Viewer with IGV_2.14 igvtools count^36,37^. Reads were trimmed and verified for quality with Trim Galore! (v.0.3.7, RRID:SCR_011847; https://www.bioinformatics.babraham.ac.uk/projects/trim_galore/). Trimmed reads were aligned with STAR (v.2.7.10a^38^, RRID:SCR_004463) to the combination of human genome and ERCC92 spike-in sequences in a single pass using default settings for gene counting (--quantMode GeneCounts). Transcripts with < 0.5 million reads mapped in all samples were removed before differential expression (DE) analysis. ERCC-invariant sequences were used with RUVg (RUVSeq v.1.44.0 ^39^) with k=1 under R (v.4.5.1 ^40^) to capture unwanted variation. The resulting weight matrix was incorporated into the DESeq2 (v.1.51.1 ^41^, RRID:SCR_000154) design matrix.

All time points (18, 24, 36, or 48 hrs) were used to establish which genes varied over time as follows: any gene that varied at any time vs its DMSO control in the same direction (up- or down-regulated) with FDR < 0.05 was deemed DE for purposes of intersection across all time points. DE genes thus identified as being consistently up- or down-regulated were then used separately in Metascape (online version, https://metascape.org^42^, RRID:SCR_016620) for pathway analysis. Heat maps were produced with the R package pheatmap (v.1.0.13 ^43^, RRID:SCR_016418) using only time course data for either 10-15 plus HCC95 (genes shown are DE for 10-15 cells) or 10326 plus HCC95 (genes shown are DE for 10326 cells). In all cases, log2 fold-changes (log2FC) vs DMSO control are shown with the row order set from largest to smallest log2FC at 48 hrs in either 10-15 or 10326 cells. No genes were called DE in HCC95 across all time points. On the left are annotation ticks showing genes in Metascape pathways (10-15 cells: up R-HSA-9012999: Rho GTPase cycle, down M66: PID MYC ACTIV PATHWAY; 10326 cells: up R-HSA-19345: Signaling by Rho GTPases, down M66: PID MYC ACTIV PATHWAY).

### Cleavage Under Targets & Release Using Nuclease (CUT&RUN)

Trypsinized 10-15, HCC95, and 10326 cells were fixed in suspension with 1% paraformaldehyde (Electron Microscopy Sciences, 15714) for 60 seconds in a 15mL conical. Fixed cells were then harvested at room temperature, resuspended in Wash Buffer (20 mM HEPES pH 7.9, 150 mM NaCl, 0.5 mM spermidine, 1× Roche protease inhibitor cocktail), and counted using a Vi-CELL Blu cell counter. For each antibody, 500,000 (HCC95) or 1 million (10326, 10-15) cells were incubated for 5-10 minutes with 10 µl of CUTANA Concanavalin A-conjugated magnetic beads (Epicypher #21-1401, Durham, NC) in Bead Activation Buffer (20 mM HEPES pH 7.9, 10 mM KCl, 1 mM CaCl2, 1 mM MnCl2). Bead-bound cells were permeabilized in 50 µL Antibody Buffer (20 mM HEPES pH 7.9, 150 mM NaCl, 0.5 mM spermidine, 0.1% digitonin , 2 mM EDTA). Samples were then incubated with 1 µL of anti-H3K27ac antibody (Cell Signaling Technologies #8173, RRID:AB_10949503), anti-BRD4L antibody (EpiCypher #13-2003, RRID: AB_3076424), anti-NUT antibody (Cell Signaling Technologies #3625, RRID:AB_2066833), or rabbit α-mouse IgG (abcam #ab46540, RRID:AB_2614925) overnight at 4°C with nutation. Bead-bound cells were washed twice with 200 µL Wash Buffer + 0.1% digitonin, and then incubated with 2.5 µL CUTANA pAG-MNase (EpiCypher, Durham, NC) in 50 µL Wash Buffer + 0.1% digitonin for 1 h at 4°C with nutation. Bead-bound cells were washed twice in 200 µL Wash Buffer + 0.1% digitonin. Cells were resuspended in 200 µL Wash Buffer (20 mM HEPES pH 7.9, 0.5 mM spermidine, 0.1% digitonin, 1x Roche protease inhibitor cocktail), the pAG-MNase activated for 30 minutes at 4°C with 1 µL 100 mM CaCl2, the reaction stopped and chromatin fragments were released by the addition of 50 µL Stop Buffer supplemented with 0.5ng E. coli spike-in DNA (340 mM NaCl, 4 mM EGTA, 20mM EDTA, 0.01% digitonin, 50 µg/ml RNase A, 50ug/mL Glycogen) followed by incubation for 30 min at 37°C. Once the pAG-MNase reaction is halted, the supernatant, containing cleaved chromatin, was incubated at 55degC overnight with 1µL of 10% SDS and 1 µL 10 mg/mL Proteinase K.

To extract the DNA, regardless of cell type, 200 µl of Oligo Binding Buffer (Zymo) followed by 800 µL of 100% ethanol was added to each sample and the total volume was then loaded onto a Zymo-Spin DCC-5 by centrifugation. The column was washed twice with 200 µL Zymo DNA Wash Buffer and then the DNA was eluted from the column in 15 µl of DNA Elution Buffer. Sample concentration was measured using a Qubit High Sensitivity dsDNA Assay on a Qubit 2.0 Fluorometer (Invitrogen). Illumina libraries were prepared using a NEBNext Ultra II Library Prep Kit (New England Biolabs) for Illumina with NEBNext Multiplex Oligos for Illumina with the following modifications. All volumes were reduced 2-fold. The NEBNext Adaptor was diluted 1:25 in NEBNext Adaptor Dilution Buffer. After Adaptor Ligation, a DNA cleanup using 1.0x volumes of Sera-Mag Select beads (Cytiva #29343052) was conducted. The PCR thermal cycling conditions were as follows: 1 cycle: 98°C for 45 s, 15 cycles: 98°C for 15 s, 65°C for 10 s, 1 cycle: 72°C for 1 min, hold: 4°C. After PCR amplification, DNA cleanup was conducted twice with 1.0x volumes of Sera-Mag Select beads. Quality control was conducted on the resulting 15 μl eluate using an Agilent 4200 TapeStation D1000 ScreenTape to determine sample concentration and sample quality. Libraries were sequenced on an Illumina NovaSeqX Plus to generate 50 bp paired-end reads.

### Analysis of CUT&RUN sequencing

Immunoprecipitations (IP, n=2 in all cases) were: IgG (negative control), H3K27ac (positive control and megadomain determination), BRD4-NUT (megadomain determination except for HCC95 with does not express BRD4-NUT), BRD4long (wild type BRD4 protein). All samples were processed under a conda environment (https://conda.org) to ensure consistent package usage under Ubuntu 22.04.2 LTS (GNU/Linux 5.19.0-1026-aws x86_64). The paired raw sequence files should be in “sample/_qc_output/_fastqc/_output/_debug”. A genome combining the Ensembl human genome as above in “RNA Sequencing” with E. coli MG1655 spike-in sequences downloaded from the NCBI (https://www.ncbi.nlm.nih.gov/datasets/genome/GCF_000005845.2/) was used for QC and QC alignment. Initial quality control (QC) was done using BWA (v.0.7.17^44^, RRID:SCR_010910). Adapters were trimmed using trimmomatic PE (v.0.36, ^45^, RRID:SCR_011848) with TruSeq3-PE adapters. Final alignment was to the same combined genome but used bowtie2 (v.2.3 ^46^, RRID:SCR_016368). After filtering on paired (-F 4 -f 2), good quality (-q 30) reads, samples were deduplicated using picard MarkDuplicates (v.2.9, https://broadinstitute.github.io/picard). After deduplication, human and E. coli sequences (divided by 2 since both ends were counted) were calculated with samtools view^47^ (RRID:SCR_002105). So-called “green list”^48^ were counted using deepTools multiBamSummary BED-file (v.3.4 ^49^, RRID:SCR_016366) for each file and cell type to normalize canonical human chromosomes (chr1-chr22, chrX, chrY) using deepTools bamCoverage with the “scaleFactor” calculated from the green list for each sequence file. These files were then used for later domain counts and signal strength as they were already normalized. Epic2 (v.0.0.52^50^) was used for peak finding with deduplicated bam files (not normalized files) as the controls provide for background frequency to identify peaks. Each sample was run against its cognate control input (e.g., for 10-15 cells, the H3K27ac, BRD4-NUT, and BRD4long IPs were all run agains their cognate 10-15 IgG input). Peaks were then used as follows: each peak per technical repeat was extended to 5Kb and merged individually using bedTools mergeBed (v.2.30 ^51^, RRID:SCR_006646); replicates were then merged without further extension; peaks that were present in both replicates were retained for further domain assessment.

For each cell line, regions of interest were counted on normalized green bigwig (.bw) files based on the bed files for each region, whether before (pre) or after (post) domain assessment, yielding both IP and IgG counts per replicate. For figures containing summarized counts, individual bigwig files were merged using bigWigMerge and bedGraphToBigWig scripts from UCSC (https://hgdownload.soe.ucsc.edu/admin/exe/linux.x86_64). For cells containing BRD4-NUT (all but HCC95), megadomains (MD) are defined as (a) above the inflection point (as defined by ROSE ^52^) for H3K27ac based on either size of region or signal over the region and (b) also BRD4-NUT positive by overlap using “bedTools intersectionBed” for H3K27ac and BRD4-NUT in regions defined above in peak mergers. After removing MD, super-enhancers (SE) lie above the H3K27ac inflection point but do not contain BRD4-NUT. The remaining peaks, so-called “regular” domains, all lie below the H3K27ac inflection point.

### CRISPR-Cas9 screen reagents and procedures

The CRISPR screen was conducted in biological duplicate using the Brunello (CP0041)^35^ libraries purchased from the Genetic Perturbation Platform at The Broad Institute (RRID:SCR_007073). 10-15-Cas9 cells were infected with pooled CRISPR library virus using polybrene (8 µg/ml, Sigma) using a low multiplicity of infection (MOI = 0.75) to achieve an average representation of ∼500 cells per gRNA for the Brunello virus. The transduced 10-15 bCas9 cells were selected using 1ug/mL puromycin (Clontech) for five days. After harvesting a subset of cells on day zero, the remaining cells were divided into two groups and treated with DMSO or 750 nM DC-9476 for 14 days, and were then harvested using Machery-Nigel Nucleospin Blood gDNA extraction kits. Bar-coded plasmid DNA was amplified via PCR and sent for next-generation sequencing (NGS) at the Broad Institute. Quantification of NGS data was performed using PoolQ (https://portals.broadinstitute.org/gpp/public/software/poolq), and Apron v1.0 (https://portals.broadinstitute.org/gppx/apron2/screener) was used to rank the performance of individual genes based on enrichment comparing the DC-9476 treatment group with the DMSO-treated group or library plasmid DNA (pDNA). Log2-transformed normalized read counts were scored for gene effect by averaging the fold change for constructs sharing the same target relative to either pDNA or DMSO read counts. A z-score was then assigned to each gRNA construct and to each target gene, the latter based on average log (fold change) relative to pDNA or DMSO samples. gRNA constructs with low abundance (z ≤ −3) in the DMSO condition were excluded from analysis. Statistical significance per target was calculated from target gene z-scores using the standard distribution of the average log(fold changes). Average log(fold change) and −log10(p-value) were plotted for genes with three or greater constructs remaining.

### Cell Cycle Analysis by Flow Cytometry

10×10^^6^ cells were trypsinized and spun at 500 x g for 5 minutes at 4 °C, and washed twice with PBS. Pellets were resuspended in 200µL of cold PBS and added dropwise while gently vortexing to 800µL 100 % ethanol in a 2mL Eppendorf tube. Fixed cells were then frozen at –20 °C for a minimum of 2 hours. Prior to staining,, cells were centrifuged at 500 x g for 5 minutes at 4 °C and washed twice with 1 ml of cold PBS. Cells were resuspended in 500 µl of propidium iodide (PI) staining solution (0.2 mg/ml RNAse A, 0.05 mg/ml PI, 0.1 % Triton-X in PBS) and incubated for 20 minutes at 37 °C. Samples were then transferred to ice and analyzed on a BD Symphony A3 flow cytometer at the Harvard Medical School flow cytometry core (https://immunology.hms.harvard.edu/resources/flow-cytometry). Cell cycle analysis was performed using FlowJo version10.10.0 (FlowJo, Ashland, OR, RRID:SCR_008520).

### AnnexinV-FITC Viability Analysis by Flow Cytometry

5×10^6^ cells were trypsinized and spun at 500 x g for 5 minutes at 4°C and washed twice with PBS. Pellets were resuspended in 1mL of cold 1X Binding Buffer and promptly placed on ice for the remainder of the procedure. 200µL of each sample was then subjected to staining with BD Pharmingen™ FITC Annexin V Apoptosis Detection Kit I (RRID: AB_2869082) with 10µL of AnnexinV-FITC conjugated antibody and 2µL of PI counterstain. Samples incubated at RT for 15 minutes shielded from light. After incubation, samples were diluted with 300µL of 1X Binding Buffer, transferred to ice, and analyzed on BD Symphony A3 flow cytometer at the Harvard Medical School flow cytometry core (https://immunology.hms.harvard.edu/resources/flow-cytometry). Cell cycle analysis was performed using FlowJo version10.10.0 (FlowJo, Ashland, OR, RRID:SCR_008520).

### Xenograft efficacy and pharmacodynamic studies

All *in vivo* studies were conducted at Dana-Farber Cancer Institute with the approval of the Institutional Animal Care and Use Committee in an AAALAC accredited vivarium. NMC models were established by injecting 1 x 10^6^ PER403 or 10-15 cells cells in 7-week old female NSG mice obtained from Jackson Laboratory (Bar Harbor, ME, RRID:IMSR_JAX:005557). Bioluminescent imaging was performed to verify disseminated tumor establishment in mice by injecting D-luciferin subcutaneously at 75 mg/kg (Promega) and imaged with the IVIS Spectrum Imaging System (Perkin Elmer). To quantify bioluminescence, identical regions of interest were drawn and the integrated total flux of photons (the sum of the prone and supine values) using the Living Image software (Perkin Elmer) were used for initial randomization of mice into various treatment groups and subsequently weekly imaging for assessing tumor response. Mice implanted with PER403 and 10-15 cells were randomized 7 days after cell implantation and disseminated tumor was established based on BLI signal exceeded 1.5×10^6^.

The PDX tumors, R25-02 and STG-298 were derived from a surgical samples, and implanted directly into the subcutaneous space in female NSG mice. Following initial implantation, the models were expanded and passaged continually as subcutaneous tumors in mice. For the efficacy studies, tumor fragments were dipped in Matrigel and implanted subcutaneously in 7-week old female NSG mice. Tumors were allowed to establish to an average tumor volume of ∼90 mm^3^ before randomization into various treatment groups.

DC-9476 was formulated in 5% DMSO, 50% PEG 400 and 45% of 20% 2-hydroxypropyl-ß-cyclodextrin in water and administered twice daily by oral gavage for 28 days. ABBV-075 and ABBV-744 compounds were formulated in 2% DMSO, 30% PEG 400, and 68% Phosal-50PG, and administered once daily by oral gavage for 28 days. Taz was formulated in 0.5% Methylcellulose (400cP) + 0.1% Tween 80 in water, pH4.0, and administered by oral gavage twice daily for 28 days. Vehicle-only treatments (2% DMSO, 30% PEG 400, and 68% Phosal-50PG) were administered once daily by oral gavage. Bioluminescent imaging was performed once weekly after treatment initiation and body weights were measured twice weekly.

For pharmacodynamic (PD) studies, tumor bearing NSG mice were treated with either vehicle control (2% DMSO, 30% PEG 400, 68% Phosal-50PG) or ABBV-744 (37.5 mg/kg, once daily administered for 5 days. After the last dose, three animals per group were euthanized at 2 hours and tumor samples (e.g. ovaries, liver, brain) were collected and fixed in 10% buffered formalin for immunohistochemistry.

To compare survival statistically, log-rank (Mantel-Cox) test was used.

### Accession numbers

RNA-seq data will be available in Gene Expression Omnibus (GEO; RRID:SCR_005012) with accession number GSE332774. CUT&RUN data will be available in GEO with accession number GSE332674.

## Results

### Discovery of a BRD4-, BD2-selective bromodomain inhibitor, DC-9476

To evaluate the activity and selectivity of BD2 inhibition in NC, we compared the BD1-BD2-BETi ABBV-075 with the BD2-selective BET inhibitor ABBV-744 across NC and non-NC cell lines, incorporating previously reported NC data^18^ together with newly generated data (**Fig. 1A**). This expanded analysis shows that NC cells are highly sensitive to BD2i, with potency comparable to BD1-BD2-BETi, whereas non-NC cells, including 293T and the lung squamous carcinoma cell lines Calu-1, HCC95, and H520, show variable sensitivity to ABBV-075 but little to no response to ABBV-744 (**Fig. 1A**). Together with the improved tolerability of BD2-selective BET inhibitors^27,29,30^, these findings support a broadened therapeutic window for NC.

**Fig. 1.**
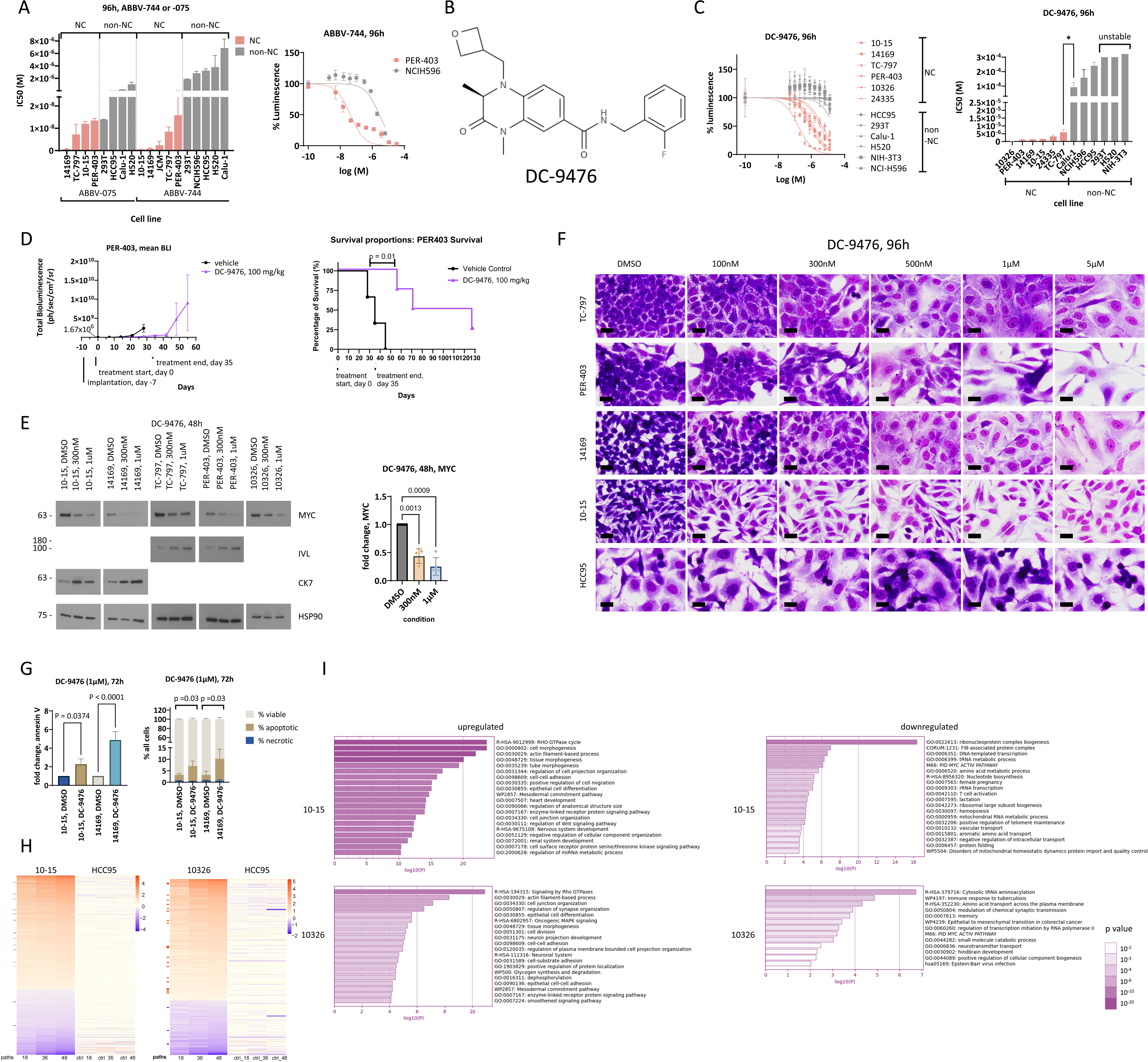
DC-9476, a novel BRD4-selective BD2i, is a potent, selective inhibitor of NC**. A.** (Left) Comparison of IC50s of ABBV-744 and ABBV-075 in NC and non-NC cell lines. Some NC cell-line data were previously reported in^18^; the graph is adapted from^18^ to include additional NC and non-NC cell lines. 14169, JCM, TC-797, 10-15, and PER-403 are BRD4-NUT+ NC cell lines. Calu-1, HCC95, and H520 are non-NC lung squamous, or adenosquamous (NCIH596) carcinoma cell lines. (Right) Dose response curve of NCIH596, a non-NC lung non-small cell carcinoma cell line identified in DepMap as the second-most sensitive non-NC cell line to ABBV-744. It is 176-fold less sensitive to ABBV-744 than the least sensitive NC cell line, PER-403. **B.** Structure of DC-9476. **C.** (Left) Dose-response curves using CellTiter-Glo as readout of cell growth/viability. 24335, ZNF532-NUT+ NC cell line; 10326, BRD3-NUT+ NC cell line. (Right) Corresponding IC50 values based on triplicate experiments. **D.** (Left) Growth of PER-403 disseminated xenograft model treated as indicated. (Right) Corresponding Kaplan-Meier survival plots. **E.** (Left) Immunoblots using antibodies to the proteins as indicated. IVL, involucrin (marker of terminal squamous differentiation. (Right) Corresponding Repeated-measures one-way ANOVA (RM one-way ANOVA) analysis. **F.** Representative microscopic images of cells grown on coversilp treated as indicated and stained by Hemacolor (MilliporeSigma). Scale bar, 20µm. **G.** (Left) Cells treated as indicated in biologic triplicate were analyzed for apoptosis and necrosis by flow cytometric analysis of Annexin V and propidium iodide (PI) staining and analyzed by RM one-way ANOVA. (Right) Percent viable, apoptotic, and necrotic cells calculated based on experiment on left. Comparisons of percent necrotic plus apoptotic cells were made by paired t-tests with p-value indicated. **H.** Heat maps of RNAseq including the superset of differentially expressed (DE) genes of DC-9476-treated cells compared to DMSO based on all time points (18h, 24h, 36h, 48h) with a FDR cutoff < 0.05 showing any change in the same direction (up or down). Heatmap from 24h time point not shown. Samples were taken from biologic duplicates. **I.** Metascape heatmaps depicting enriched Genome Ontology (GO), Reactome Homo sapiens (R-HSA), WikiPathways (WP), Comprehensive Resource of Mammalian Protein Complexes (CORUM), Pathway Interaction Database MYC Activation Pathway (M66), and Homo sapiens (hsa) gene sets compared with DMSO-treated cells identified by GSEA for DC-9476 treatment based on RNAseq corresponding to **H.**

Because all of our non-NC cell lines were insensitive to ABBV-744, we interrogated the Cancer Dependency Map Project at Broad Institute (DepMap, https://depmap.org) to identify ABBV-744-sensitive non-NC cancer cell lines. The two most sensitive cell lines of 888 tested to ABBV-744 (2µM) are lung non-small cell carcinomas, RERFLCKJ and NCIH596, with log2 (fold change) cell abundance relative to vehicle control of −8.3 (99.99 percentile) and −7.4 (99.8 percentile), respectively (mean for all cell lines, −1.43) (**Supplementary Fig. S1A**). We acquired and treated NCIH596 cells with ABBV-744, revealing that this ABBV-744-“sensitive” non-NC cell line, while sensitive to high micro-molar doses, is no more sensitive than 293T cells and is 176-fold less sensitive than our least ABBV-744-sensitive NC cell line, PER-403 (**Fig.1A**).

Our findings that BD2i is uniquely active in NC in vitro prompted us to investigate whether an even more selective, and thus better tolerated, BD2i may also inhibit NC growth. Given that the majority of NCs harbor *BRD4::NUTM1* fusions, we sought to design and synthesize a BRD4-, BD2-selective inhibitor.

Using DeepCure’s integrated physics-based artificial intelligence (AI) platform, we developed a multi-stage computational design workflow. First, PocketExpander™ combined molecular dynamics with quantum physics-based modeling to identify transient binding opportunities unique to BRD4-BD2^53^. MolGen™, a distributed 3D deep reinforcement learning platform, then generated and optimized candidate ligands, guided by SAPT-inspired Equivariant Graph Neural Networks that capture key intermolecular interactions. Finally, DeepEnergy™ then prioritized molecules by evaluating interaction stability and developability across molecular dynamics trajectories ^54–56^.

This workflow produced a novel 3,4-dihydroquinoxalin-2(1H)-one scaffold with higher affinity for BD2 over BD1 and modest selectivity for BRD4-BD2. Subsequent optimization of potency, selectivity, and absorption, distribution, metabolism, and excretion (ADME) properties yielded DC-9476 (**Fig. 1B**), a lead compound suitable for clinical development ^57^. In time-resolved fluorescence resonance energy transfer (TR-FRET) assays, DC-9476 showed 1,440-fold higher affinity for BRD4-BD2 over BD1, and 9.1- and 22.9-fold higher affinity for BRD4-BD2 over BRD3-BD2 and BRD2-BD2, respectively (**Table 1**). These findings establish DC-9476 as highly selective for BD2, with modest selectivity for BRD4.

### NC is uniquely sensitive to the BRD4 -BD2-selective inhibitor, DC-9476, in vitro and in vivo, regardless of the NUT-fusion partner

We next sought to determine whether DC-9476 can inhibit NC cells. Predictably, this compound was nearly completely inactive in six non-NC cell lines, including two non-neoplastic and four lung squamous cancer cell lines, including the ABBV-744 “sensitive”, NCIH596 cells (**Fig. 1C**). In contrast, all of six NC cell lines were uniquely sensitive to DC-9476, showing substantial growth repression with IC50s of 144nM-5.6µM based on CellTiter-Glo viability assays (**Fig. 1C**). Unexpectedly, the NC cell line most sensitive to BRD4-BD2 inhibition by DC-9476, 10326, harbors a *BRD3-NUTM1* fusion. Also unexpectedly, the ZNF532-NUT+ NC line, 24335, was even more sensitive to DC-9476 than one of our BRD4-NUT+ cell lines, TC-797. These findings raise the possibility that at least part of the inhibitory activity of DC-9476 may be through inhibition of wild type BRD4, potentially uncovering an unexplored role of this protein in BRD-NUT pathogenesis. However, the higher doses of DC-9476 used in these assays exceeds that predicted to be selective for BRD4-selective inhibition, thus necessitating further evaluation.

The in vitro potency of DC-9476 motivated us to evaluate its activity in an in vivo preclinical model of NC. To model NC, we inject, via the tail vein, NOD-SCID-GAMMA (NSG) mice with NC cell lines that express luciferase^58^. Upon injection these cells disseminate to solid organs, such as ovary and liver, and bone, within one week, robustly recapitulating the human disease. After disseminated tumor was established based on bioluminescence (BLI) signal ≥ 1.5×10^6^ ph/sec/cm²/sr, we administered vehicle control or DC-9476 (100mg/kg twice daily) for 35 days, after which treatment was terminated and mice were observed for up to 130 days.

In the PER-403 BRD4-NUT+ xenograft model (n = 4 mice per arm), we found that DC-9476 treatment was well tolerated (**Supplementary Fig. S1B**), leading to durable growth inhibition and significantly prolonged survival (**Fig. 1D**). These findings indicate that DC-9476 may be an effective, well-tolerated treatment for NC either alone or combined with synergistic targeted inhibitors.

### BRD4/BRD3-BD2-selective inhibition phenocopies the effects of BRD-NUT depletion

Taking advantage of the unique specificity of DC-9476, we utilized this molecule to determine how selective inhibition of BD2 of BRD4 and, to less of an extent, BRD3, affects growth and differentiation. Thus, NC cells were treated with doses of DC-9476 sufficiently low to inhibit BD2 of BRD4, BRD3, or BRD4/BRD3-NUT (300nM), and not other BET proteins. A non-NC squamous lung cancer cell line, HCC95, was also treated with these doses. We observed that these doses of DC-9476 treatment caused rapid loss of MYC (**Fig. 1E**) in all NC cells, but not in HCC95 cells (**Supplementary Fig. S1C**), a finding consistent with inhibition of BRD-NUTs function to maintain MYC expression^16,17^. This finding was observed both in BRD4-(10-15, 14169, PER-403, TC-797) and BRD3-NUT+ (10326) cell lines.

MYC loss in BRD4-NUT+ NC cells treated with DC-9476 corresponds with robust differentiation, evidenced by upregulation of squamous (involucrin, IVL) or epithelial (CK7) markers (**Fig. 1E**), and dose-dependent morphologic changes of squamous differentiation consisting of flattening and enlargement of cells (**Fig. 1F**). Accompanying differentiation is significant induction of apoptosis and necrosis, as determined by flow cytometric analysis of Annexin V and propidium iodide (PI) staining (**Fig. 1G**), suggesting that therapeutic treatment with DC-9476 can both block NC tumor growth and induced cell death. Features of differentiation can overlap with senescence, however we found that senescence was not a consistent effect of DC-9476, with some NC cell lines showing a trending increase of beta-galactosidase staining, and others showing a decrease (**Supplementary Fig. S1D**).

We next determined what transcriptional changes occur in response to inhibition by DC-9476 by transcriptomic analysis of a BRD4-NUT+ (10-15) and BRD3-NUT+ (10326) NC cell line, and in non-NC HCC95 cells. We performed ribosomal-RNA-depleted RNA sequencing (RNAseq) to comprehensively identify differentially expressed (DE) genes in response to DC-9476 treatment using a BRD4-/BRD3-, BD2-selective dose (300nM). Samples were collected at four time points: 18h, 24h, 36h, and 48h. To simplify analysis, we compiled a superset of DE genes from all time points with FDR cutoff < 0.05 showing any change in the same direction (up or down). Remarkably, over the time course of treatment, no such DE genes were identified in HCC95 cells. In contrast, 315 and 64 genes were differentially downregulated, and 550 and 142 genes were differentially upregulated in 10-15 and 10326 cells, respectively (**Fig. 1H, Supplementary Fig. S2**). Gene set enrichment analysis (GSEA) using Metascape identified consistently upregulated and downregulated pathways in both NC cell lines. Pathways associated with epithelial differentiation (i.e., RHO GTPase, cell-cell adhesion, epithelial differentiation) were significantly enriched with differentially upregulated genes in treated 10-15 and 10326 cells (**Fig. 1I, Supplementary Tables S6-9**). Conversely, downregulation of MYC and protein translation pathways were observed in both treated 10-15 and 10326 cells. Overall, the changes in expression induced by DC-9476 match what is observed phenotypically and that which has previously been reported for BD1-BD2-BETi^8,12,17,18,20,32,59,60^: MYC and MYC pathway inhibition, epithelial differentiation and growth arrest in NC cells, and lack of response in non-NC cells. These transcriptional-phenotypic effects are specific to the function of BRD4 and BRD4-NUT to block differentiation through upregulation of MYC^16,17^.

Taken together, the phenotypic and transcriptomic responses to inhibition by DC-9476 are consistent with on-target inhibition of BRD-NUT in NC, thus identifying DC-9476 as a potent, selective inhibitor of NC.

### BD2 inhibition induces proteasomal degradation of BRD4 and BRD4-NUT in vitro and in vivo

Previous studies have shown that various factors, including cell cycle components (i.e., EZH2-CDK4/6-Rb axis), confer resistance to BD1-BD2-BETi^18,61^ in NC. We thus employed a CRISPR knockout screen to identify unique factors that might mediate sensitivity to BD2i. In this screen, 10-15 NC cells infected with the Brunello whole genome sgRNA library^35^ were treated with DC-9476 (750nM) or DMSO for fourteen days to identify genes whose loss leads to increased sensitivity or resistance to BD2i. By a large margin, the two top resistance hits were Speckle Type BTB/POZ Protein (SPOP), an E3 ligase that targets BET proteins, including BRD4, for degradation^62^, and Cullin3 (CUL3), a key core component of the E3 ubiquitin ligase complex^63^ containing SPOP^64^ (**Fig. 2A, Supplementary Table S10**). SPOP inactivating mutations impart resistance to BD1-BD2-BETi through impaired degradation of, and thus accumulation of BRD4^62^. While these studies provide a mutational strategy for the development of cancer resistance to BETi, they rely entirely on the assumption that altered BET protein stoichiometry enables BET proteins to outcompete BETi for chromatin binding.

**Fig. 2.**
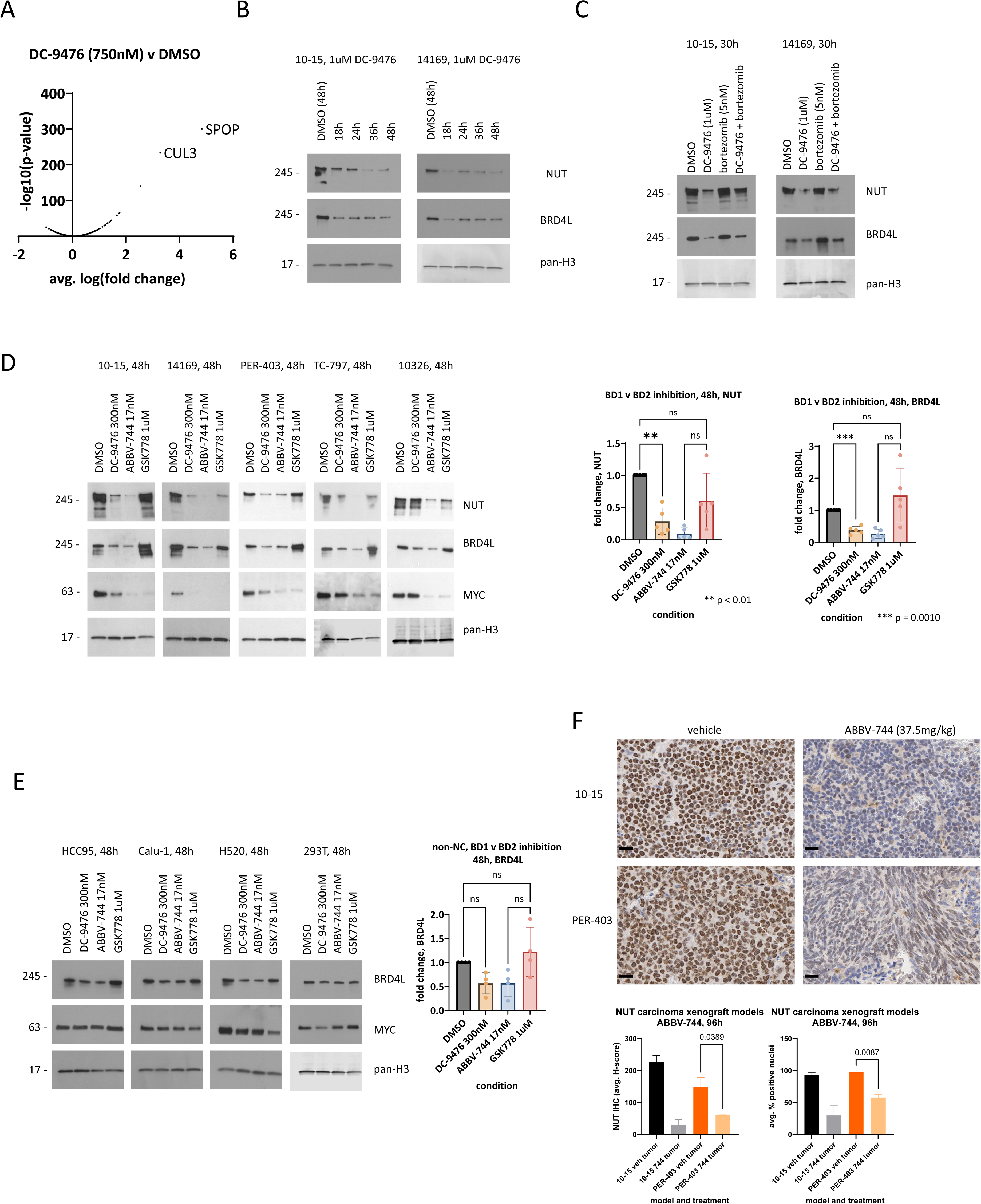
BD2 inhibition induces proteasomal degradation of BRD4 and BRD4-NUT in vitro and in vivo. **A.** Plot of log (fold change) averaged for each gene vs. p-values from the CRISPR knockout screen comparing DC-9476-with DMSO-treated 10-15-Cas9 cells. Shown is a representative single biological replicate from duplicate experiments. **B.** Immunoblot of NC cells treated over a time course as indicated. **C.** Immunoblot of NC cells treated as indicated. Doses of each compound used in combination are the same as each used as single agents. **D.** (Left) Immunoblot of NC cells treated as indicated. DC-9476 and ABBV-744 are selective BD2i; GSK778 is a selective BD1i. (Right) Corresponding RM one-way ANOVA analysis. **E.** (Left) Immunoblot of non-NC cells as indicated. (Right) Corresponding RM one-way ANOVA analysis of group band densities at left. **F**. (Left) Photomicrographs of anti-NUT immunohistochemical (IHC) staining of formalin-fixed, paraffin-embedded sections of representative tumors explanted from mice 96h following treatment as indicated. Scale bars = 20µm. (Right) Paired t-test analysis of corresponding NUT IHC by QuPath (n = 3 mice per arm) using two scoring systems – H-score and % positive nuclei.

The striking dependence of BD2i resistance in NC on SPOP-CUL3, with both encoded by genes that are dominant outlier hits in our CRISPR screen, suggests that resistance is not explained solely by altered BET protein stoichiometry, but may instead reflect protection of BRD4 and BRD4-NUT from BD2i-induced loss. Indeed, treatment of NC cell lines with DC-9476 leads to depletion of both wild type BRD4 (specifically the long isoform of wild type BRD4, here termed BRD4L) and BRD4-NUT within 18h (**Fig. 2B**). BRD4L and BRD4-NUT loss is at least partially due to proteasome-mediated degradation, as determined by rescue with bortezomib, a proteasome inhibitor (**Fig. 2C**). Unexpectedly, BD2 inhibition, but not BD1 inhibition, significantly depleted BRD4L and BRD4-NUT, as tested using selective inhibitors of either bromodomain (**Fig. 2D**). BD2i compounds included DC-9476 and ABBV-744; BD1i included GSK778^27^. Similarly, in non-NC cells, depletion of BRD4L by BD2i, but not BD1i, was also observed (**Fig. 2E**). Unlike NC cells, however, the degree of depletion in non-NC cells was not statistically significant, and substantial BRD4L remained after BD2i (**Fig. 2E**). Because BD1-BD2 BETi also targets BD2, we expected that it would phenocopy the effects of BD2i in NC cells. However, treatment with diverse BD1-BD2-BETi, including ABBV-075, JQ1, and OTX-015, depleted only BRD4-NUT and not BRD4L (**Supplementary Fig. S3**). These findings suggest that simultaneous BD1 and BD2 engagement may attenuate BD2i-induced degradation of BRD4L, whereas BRD4-NUT remains susceptible to degradation, perhaps by secondary events downstream of BD1-BD2 BETi.

Depletion of BRD4-NUT in NC cells by BD2i is also observed in vivo. Treatment of two of our bioluminescent xenograft models of NC, 10-15 and PER-403, with ABBV-744 for 96h resulted in loss of BRD4-NUT in explanted tumors stained for NUT immunohistochemically (**Fig. 2F**). In one mouse model (PER-403), the decrease in NUT staining was significant comparing H-scores (unpaired t-test, p = 0.0389) or percentage positive nuclei (p = 0.0087) with vehicle. In the other (10-15), ABBV-744 markedly reduced the H-score compared with vehicle (mean 33 vs 223; p = 0.056), however, this comparison is more significant than the numbers show due to complete tumor regression.

Together, these findings suggest that the selective loss of BRD4L and BRD4-NUT after BD2 inhibition -but not BD1 inhibition – may at least partly explain the unusual sensitivity of NC to BD2-selective inhibitors, with important therapeutic implications.

### BRD4L is required for NC growth and the blockade of differentiation

BRD4-NUT is the sole oncogene driver of BRD4-NUT+ NC, as established in genetically engineered mouse models of NC^12,13^. Loss of BRD-NUT or NSD3-NUT leads to rapid squamous differentiation and growth arrest^7,8^, thus its degradation induced by BD2i is expected to be catastrophic to NC cells, explaining in part the potency of this compound class in NC. However, it is also known that BRD-NUT oncogenic function is dependent on cooperation with various wild type factors, including EZH2 and MYC^17,18^. There is some evidence that BRD4L may, when phosphorylated by CDK9, be required for NC growth as well^65^. Thus, the loss of BRD4L upon BD2i treatment prompted us to test what effect BRD4L depletion has on NC cells.

Our approach was to determine the effect of CRISPR-Cas9-knockout of *BRD4L* on NC cell growth and differentiation. To attain adequate knockout of *BRD4L*, we utilized a doxycycline-inducible guide-only plasmid (pRDA355, Addgene #187159^66^) to infect our 10-15 Cas9+ cells (multiple siRNAs tested failed to adequately knock down BRD4L). Using this system, knockout of BRD4L resulted in rapid phenotypic changes consistent with epithelial differentiation and/or senescence, evidenced by cell flattening, enlargement, reduction in MYC, and increased expression of the epithelial differentiation marker, CK7, and the squamous differentiation marker, FOSL2^67^ (**Fig. 3A-B**). These changes are similar to the effects of BRD4-NUT knockdown^7^, and correspond with G1 arrest and a significant decrease of cells in S phase (**Fig. 3C-D**).

**Fig. 3.**
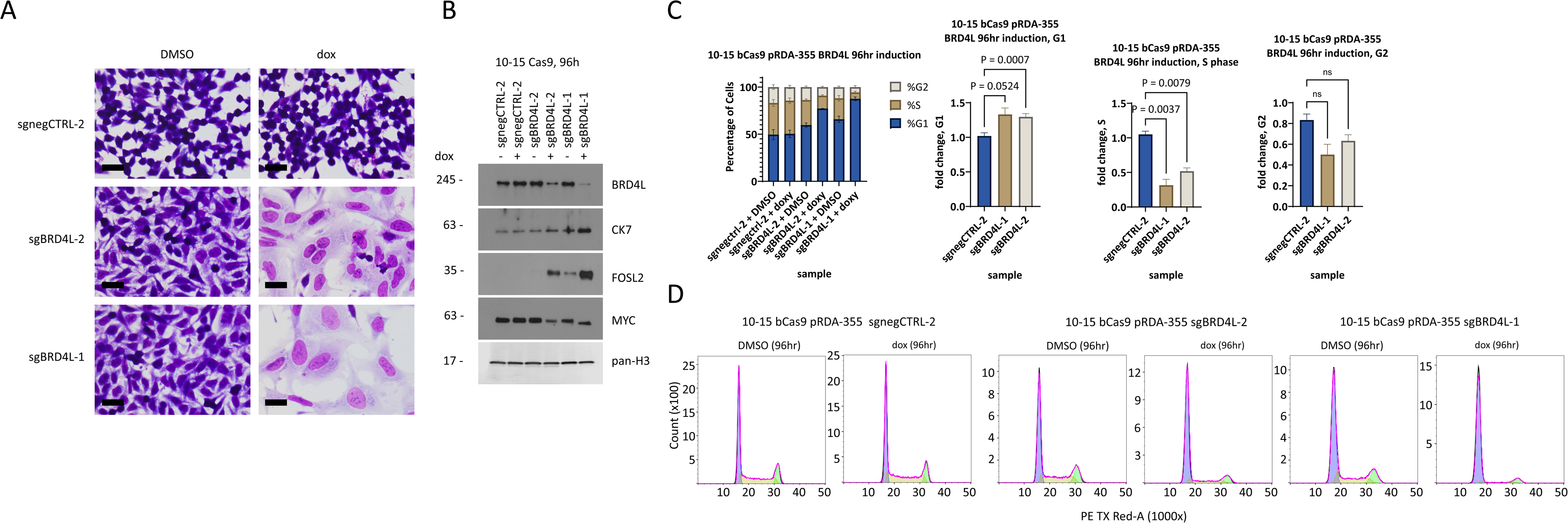
BRD4L is required for NC growth and the blockade of differentiation. **A.** Representative microscopic images of 10-15 Cas9 clones infected with pRDA355 plasmid constructs as indicated and stained by Hemacolor. sgRNA expression was induced for 96h with doxycycline (dox), or not induced using vehicle (DMSO). Scale bars = 20µm. **B.** Immunoblot of 10-15 Cas9 clones infected with pRDA355 plasmid constructs as indicated. **C.** Cells treated as indicated in biologic triplicate were subjected to flow cytometric analysis to identify proportions of cells in each phase of the cell cycle as shown and analyzed by RM one-way ANOVA. **D.** Histograms of representative single replicates corresponding to cell cycle analysis in **E**.

The findings indicate that, like EZH2 and MYC, BRD4L is a key co-factor that with BRD4-NUT is required for NC growth and to prevent differentiation. Importantly, they indicate that loss of BRD4L is likely to contribute to the anti-growth effects of BD2i in NC.

### BD2 inhibition induces SPOP-mediated proteasomal degradation of BRD4L and BRD4-NUT

Identification of *SPOP* and *CUL3* knockouts as top resistance hits to DC-9476 treatment (**Fig. 2A**) prompted us to test whether SPOP is responsible for the proteasomal degradation of BRD4L and BRD4-NUT in NC. Our strategy was to determine whether inducible CRISPR-Cas9-knockout of *SPOP* can prevent loss of BRD4L/BRD4-NUT in NC cells treated with BD2i (multiple siRNAs tested failed to adequately knock down SPOP). In a key finding, *SPOP* knockout prevented DC-9476-induced loss of BRD4L/BRD4-NUT and the corresponding decrease in MYC (**Fig. 4A**).

**Fig. 4.**
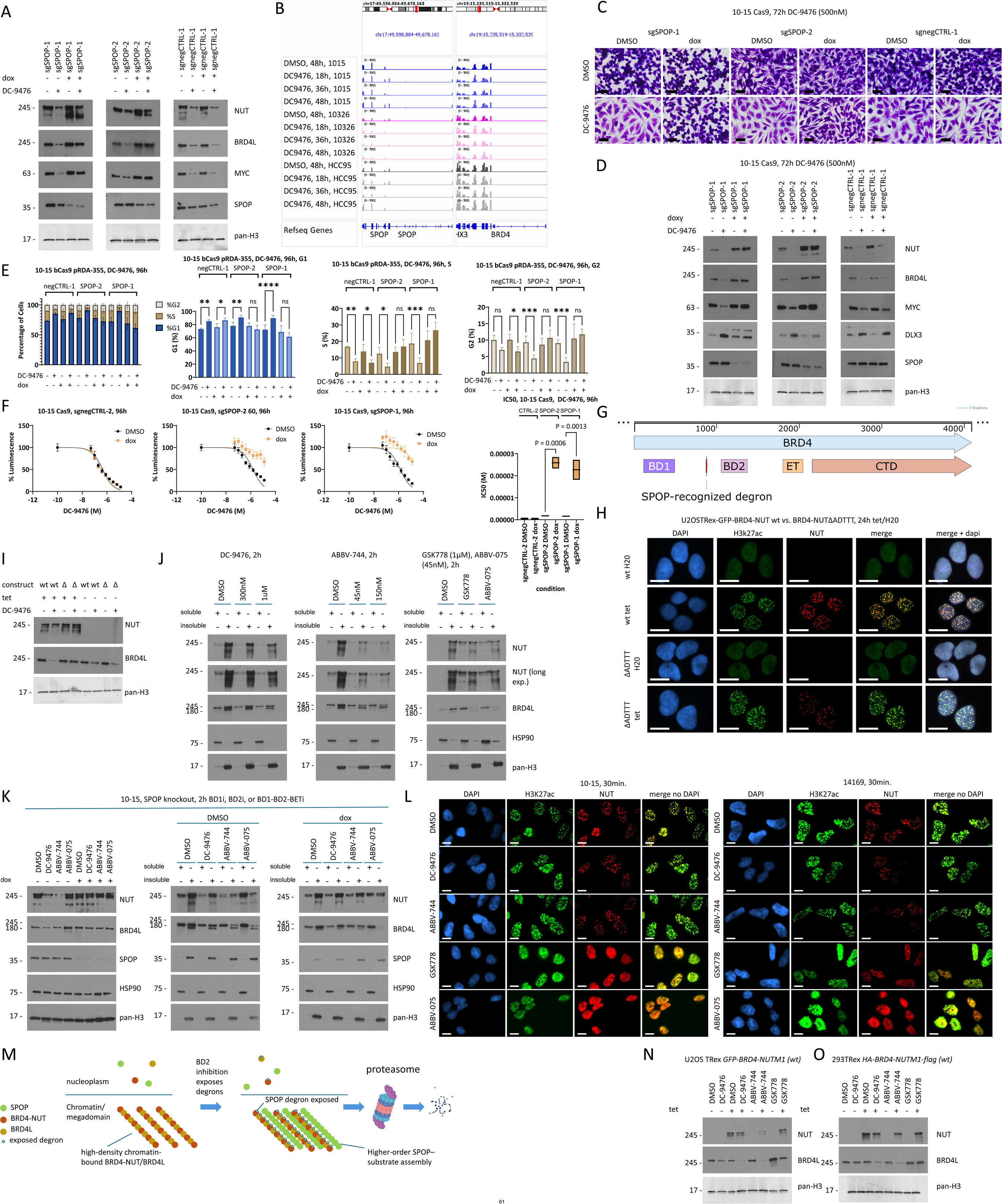
BD2 inhibition induces SPOP-mediated proteasomal degradation of BRD4L and BRD4-NUT without displacement from chromatin. **A.** Immunoblot of 10-15 Cas9 clones infected with pRDA355 plasmid constructs as indicated. DC-9476 was dosed at 300nM for 24h following 96h induction of sgRNA expression with doxycycline (dox), or not induced using vehicle (DMSO). **B.** Integrated genome viewer (IGV) views of RNAseq reads at the indicated genes. Each track shown is from one of three biologic replicates. **C.** Representative microscopic images of 10-15 Cas9 clones infected with pRDA355 plasmid constructs as indicated and stained by Hemacolor. sgRNA expression was induced in cells grown on coverslips for 96h with doxycycline, or not induced using vehicle (DMSO), followed by treatment with DMSO or DC-9476 (500nM) for 72h. Scale bars = 20µm. **D.** Immunoblot of 10-15 Cas9 clones infected with pRDA355 plasmid constructs as indicated and treated as in **C**. **E.** Cells treated as indicated in biologic triplicate were subjected to flow cytometric analysis to identify proportions of cells in each phase of the cell cycle as shown and analyzed by RM one-way ANOVA. The dose of DC-9476 was 1µM. ***\**** p <0.05**F.** (Left) Dose-response curves using CellTiter-Glo as readout of cell growth/viability. 10-15 Cas9 clones infected with pRDA355 plasmid constructs as indicated. sgRNA expression was induced for 96h, followed by treatment with DC-9476 as indicated for another 96h. (Right) Corresponding IC50 values with RM one-way ANOVA analysis. **G.** Schematic of *BRD4L* with mapped encoded protein domains and motifs as indicated. **H.** Representative immunofluorescence images of U20STRex cells induced (tetracycline (tet)) or not induced (H20) for 24h to express either wild type *GFP-BRD4-NUTM1* (wt) or *GFP-BRD4-NUTM1* with a deletion of the SPOP-recognized degron, ADTTT (ΔADTTT). Antibodies used for co-staining for NUT and H3K27ac as indicated. Scale bar, 20µm. **I.** Immunoblots of U20STRex cells induced (tet) for 9h to express *GFP-BRD4-NUTM1* (wt) or *GFP-BRD4-NUT*ΔADTTT (Δ), followed by treatment with DC-9476 (1µM) for 24h, during which tet treatment was continuous. **J.** Immunoblots of soluble and insoluble nuclear fractions purified from CKS-150 lysates of 10-15 cells treated as indicated. **K**. (Left) Immunoblots of total cell lysates from 10-15 Cas9 SPOP knockout clone SPOP-1 induced to express the sgRNA for 96h, followed by 2h treatments as indicated. (Right) Immunoblots of nuclear fractions, as described in **J**, corresponding with treatments at left. Doses are as follows: DC-9476, 1µM; ABBV-744, 45nM; ABBV-075, 45nM. **L.** Representative immunofluorescence images of BRD4-NUT+ NC cell lines, 10-15 and 14169 treated as indicated with the following doses: DC-9476, 1µM; ABBV-744, 45nM; GSK778, 1µM; ABBV-075, 45nM. Antibodies used for co-staining for NUT and H3K27ac as indicated. Scale bar, 10µm. **M.** Theoretical model of how BRD4-NUT facilitates degradation of BRD4/BRD4-NUT by SPOP. **N.** Immunoblot of U20STRex cells induced to express *GFP-BRD4-NUTM1* with tet/H20, followed by 24h treatment with DMSO, DC-9476 (1µM), ABBV-744 (17nM), or GSK778 (1µM) during which tet/H20 treatment was continuous. **O.** Immunoblot of 293TRex cells induced to express *FLAG-BRD4-NUTM1-HA* and treated as in **N**.

Given that SPOP is required for BD2i-induced degradation of BRD4/BRD4-NUT, we considered that BD2i treatment might enhance this effect by increasing SPOP expression. However, we find that SPOP RNA levels (as do those of BRD4/BRD4-NUT) remain stable in all cells tested, including 10-15, 10326, and HCC-95 cells (**Fig. 4B**). To rule out mutational gain-of-function of SPOP, as described in a subset of endometrial cancers^68^, we submitted our NC cell lines (10-15, 14169, PER-403, TC-797, 10326) for targeted next-generation sequencing (OncoPanel v3.1), but SPOP was not mutated in any of the lines (**Supplementary Table S11**).

To determine the extent to which SPOP-mediated degradation of BRD4L and BRD4-NUT contributes to DC-9476-mediated growth suppression in NC, we tested whether SPOP knockout attenuates the cellular response to DC-9476. In control sgRNA-induced cells, DC-9476 treatment led to morphologic changes characteristic of squamous differentiation, with cell enlargement and flattening (**Fig. 4C**), accompanied by upregulation of the squamous differentiation-specific transcription factor, DLX3 (**Fig. 4D**). As expected, this was accompanied by reduced MYC (**Fig. 4D**). Strikingly, induction of SPOP-targeting sgRNAs nearly completely prevented squamous differentiation and MYC downregulation induced by DC-9476 (**Fig. 4C-D**), indicating that loss of SPOP prevents suppression of the BRD4-NUT-driven transcriptional state, namely that of upregulation of MYC and its pro-growth, anti-differentiation programs (**Fig. 1I**), by BD2i. Indeed, while DC-9476 treatment leads to G1 arrest of control NC cells, consistent with what has previously been observed with BD1-BD2-BETi ^61^, those with SPOP knockout showed no changes in cell cycle (**Fig. 4E, Supplementary Fig. S4**). Together with rescue of BRD4L and BRD4-NUT protein levels, these findings demonstrate that SPOP-mediated degradation is required for the key phenotypic and transcriptional effects of BD2i. Consistent with this conclusion, SPOP knockout substantially reduced DC-9476-mediated growth suppression across a broad dose range (**Fig. 4F**), with dose-response curves resembling those of resistant non-NC cells (compare with **Fig. 1C**).

The canonical degron in BRD4 recognized by SPOPs Meprin and TRAF homology (MATH) domain, defined by the amino acid sequence, ADTTT, has previously been described^62,69^, and is localized between BD1 and BD2 (**Fig. 4G**). We thus asked whether this same degron was needed for BD2i-induced degradation of BRD4-NUT by comparing effects of BD2i on an ADTTT deletion mutant (BRD4-NUTΔADTTT) with wild type BRD4-NUT. Immunofluorescence showed that BRD4-NUTΔADTTT retained the ability to form nuclear condensates/megadomains comparable to wtBRD4-NUT, indicating that the ADTTT deletion does not grossly impair BRD4-NUT nuclear organization (**Fig. 4H**). Despite preserved foci formation, BRD4-NUTΔADTTT was resistant to BD2i-induced degradation, indicating that the ADTTT motif is specifically required for productive SPOP-CUL3–mediated degradation (**Fig. 4I**). Unexpectedly, BRD4L was also resistant to degradation, suggesting that BRD4-NUTΔADTTT dominantly protects endogenous BRD4 from BD2i-induced loss, whereas wtBRD4-NUT does not.

### BRD4L and BRD4-NUT remain chromatin-associated in the presence of BD2i

The dependence of SPOP-mediated degradation of BRD4L and BRD4-NUT on BD2i led us to hypothesize that chromatin displacement by BD2i is required for this degradation. Thus, we determined whether BD2i displaces these proteins from the insoluble, chromatin-associated fraction to the soluble fraction of nuclear extracts using CSK-based chromatin fractionation. Treatment was for only 2h to assay direct effects, and ABBV-075 was used as positive control because BD1-BD2-BETi is well documented to cause eviction of BRD4L and BRD4-NUT from chromatin ^20,70^. Unexpectedly, BD2i did not result in displacement of BRD4L/BRD4-NUT from chromatin to soluble nuclear fractions, whereas BD1i and BD1-BD2-BETi clearly evicted both proteins from chromatin to soluble fractions (**Fig. 4J**). However, overall BRD4L/BRD4-NUT abundance was reduced by BD2i. To determine whether rapid SPOP-mediated degradation might mask redistribution of BRD4 and BRD4-NUT from chromatin to nucleoplasm by BD2i, we performed a similar fractionation experiment in our inducible SPOP knockout model, 10-15 pRDA355-SPOP-1. Whole cell lysates confirmed that BD2i depletes BRD4L and BRD4-NUT within 2h, whereas BD1-BD2-BETi has no effect on protein levels; as expected, SPOP knockout prevented loss under any condition (**Fig. 4K**). In key findings, BD2i did not displace BRD4L or BRD4-NUT into the soluble fraction even in the absence of SPOP, whereas BD1-BD2-BETi did so robustly. Immunofluorescence imaging of BRD4-NUT and H3K27ac corroborated this result. At 30 min., BRD4-NUT- and H3K27ac-enriched nuclear foci corresponding with megadomains remained intact following treatment of NC cells with BD2i, DC-9476 and ABBV-744 (**Fig. 4L**). In contrast, treatment with BD1i (GSK778) or BD1-BD2-BETi (ABBV-075) led to marked diffusion of BRD4-NUT and H3K27ac staining, with loss of nuclear foci. The findings indicate that ***BD2i does not redistribute BRD4L or BRD4-NUT from chromatin-associated to soluble nucleoplasmic fractions***, a key distinguishing feature from BD1i or BD1-BD2-BETi. These findings argue against the idea that BD2i, unlike BD1i or BD1-BD2-BETi, evicts BRD4L and BRD4-NUT from chromatin. SPOP forms higher-order oligomers that are multivalent for substrate binding, and multivalent substrates can stabilize and crosslink SPOP oligomers to promote higher-order SPOP–substrate assemblies^71^. SPOP oligomerization, in turn, increases avidity for substrate and enhances CRL3^SPOP^-mediated ubiquitination^72^. BRD4-NUT-driven chromatin hyperacetylation recruits BRD4L to megadomains, resulting in a high local concentration of both proteins (**Supplementary Fig. S5**, **Fig. 5**, and^16^). We propose a model (**Fig. 4M**) whereby BD2i renders the SPOP degrons of BRD4L and BRD4-NUT accessible by changing the BD2-chromatin interface, while these proteins remain highly concentrated within megadomains, thereby generating a high-density array of SPOP-binding sites that functions as an effectively multivalent substrate platform. Because BRD4L and BRD4-NUT remain chromatin-associated following BD2 inhibition, this high local degron density is maintained, favoring SPOP oligomerization and efficient CRL3^SPOP^-mediated degradation of both proteins. Because BRD4-NUT is intrinsically less mobile on chromatin than BRD4, as shown previously^7^, retention of BRD4-NUT/BRD4L within megadomains after BD2 inhibition may further preserve a high local concentration of exposed SPOP degrons. By contrast, eviction of BRD4-NUT and BRD4L by BD1i or BD1-BD2-BETi disperses these proteins into the soluble nucleoplasmic pool, reducing their local concentration and thereby limiting efficient SPOP-mediated degradation.

**Fig. 5.**
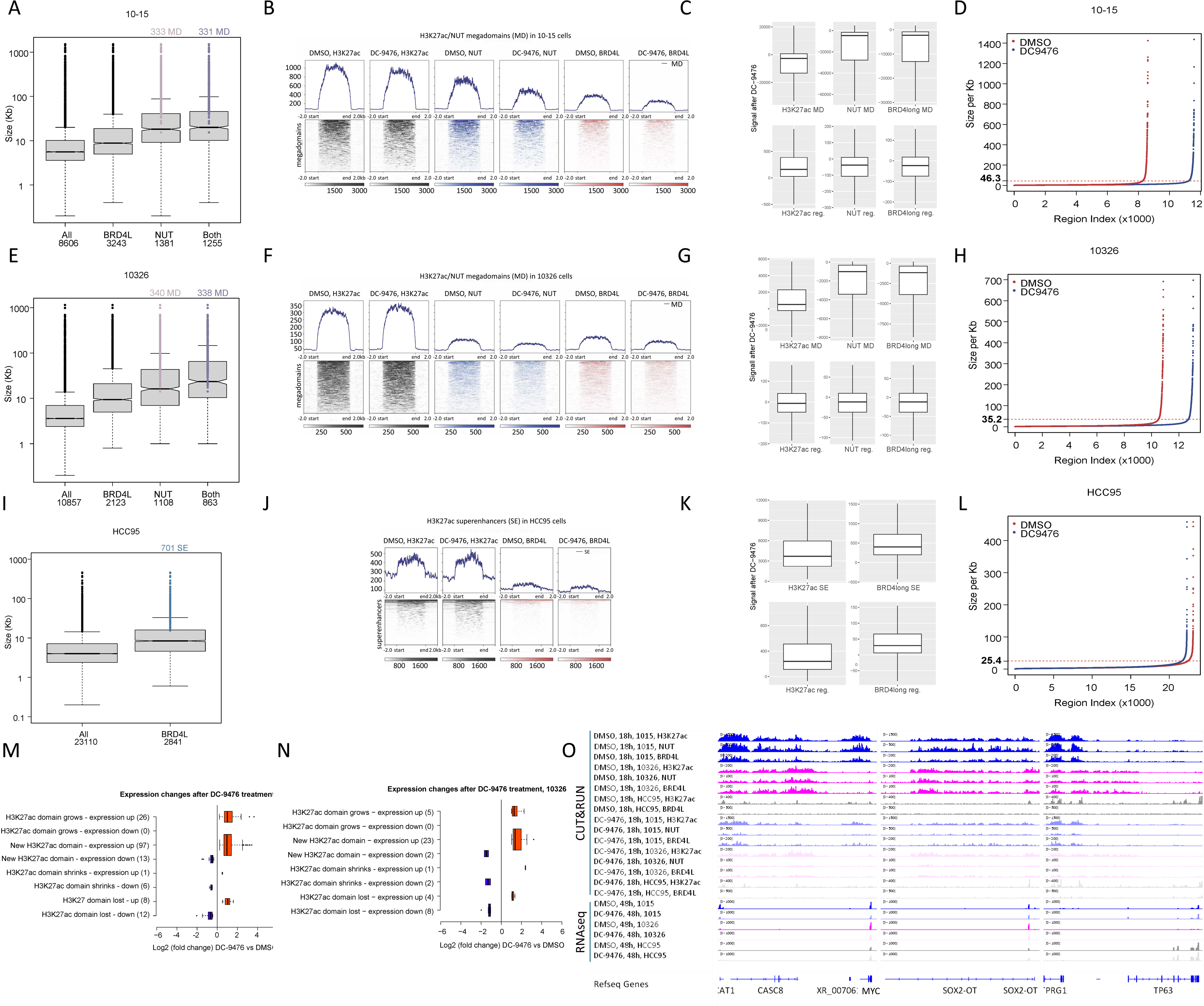
BD2 inhibition by DC-9476 depletes megadomains of BRD4L, BRD4-, and BRD3-NUT. **A.** Box plot of H3K27ac domains identified by CUT&RUN of 10-15 cells, including those containing BRD4L peaks, BRD4-NUT (NUT) peaks, BRD4L and BRD4-NUT peaks (Both), and all H3K27ac domains (All). The number of domains are indicated at bottom. The subset of NUT and “Both” domains that are megadomains (MD) are indicated at top. **B.** Profiles and corresponding heat maps of signal within megadomains identified by CUT&RUN using the indicated antibodies. Tracks are from the merger of two biological replicates. **C.** Box plots depicting the change in signal between DMSO and DC-9476, quantified by difference in reads per region (reads from DMSO-treated samples subtracted from DC-9476 treated) after treatment with DC-9476 for the indicated domains. In this plot, megadomain (MD) is defined as containing NUT and by being larger than the inflection point of the rank-ordered list by size of H3K27ac region enrichment or signal above the signal inflection point as shown in **D**. Reg. = regular enhancer. **D.** Hockey-stick plot of rank-ordered list of H3K27ac region enrichment and region size calculated from CUT&RUN performed in 10-15 cells. The horizontal dotted line indicates the inflection point above which are H3K27ac regions considered for megadomains which must also contain NUT. **E-H**, CUT&RUN data for 10326 cells as described for **A-D**, respectively. **I-L**, CUT&RUN data for HCC95 cells as described for **A-D**, respectively, except antibodies to NUT were not used due to lack of NUT expression. **M**. Correlation of transcriptional changes based on RNAseq with H3K27ac domain size changes in 10-15 cells. H3K27ac domains of all types are analyzed, including megadomains (n = 338), regular domains (n = 8,261) and super-enhancers (n = 12). Grows, domain larger after treatment; New, domain present only after treatment; Shrinks, domain smaller after treatment; Lost, domain present only in DMSO. Blue colored genes are down-regulated while red ones are up-regulated consistently in DC-9476 vs DMSO at all time points (n=4 at 18h, 24h, 36h, 48h) using FDR < 0.05 and fold-change different than 1 after 24hrs. **N**. Correlation of CUT&RUN and RNAseq as described in **M** for 10326s. H3K27ac domains include megadomains (n = 340), regular domains (n = 10,487) and super-enhancers (n = 30). **O**. Integrated genome viewer (IGV) views of CUT&RUN and RNAseq peaks at the indicated genes. Each CUT&RUN track shown is from one of two biologic replicates. Each RNAseq track shown is from one of three biological replicates.

### BRD4-NUT sensitizes BRD4L to degradation in the presence of BD2i

Our model predicts that megadomains formed by BRD4-NUT in any cell type would sensitize itself and BRD4L to BD2i-induced degradation due to the formation of highly concentrated SPOP substrate and would explain the lack of such sensitivity in non-NC cells lacking BRD4-NUT (**Fig. 2E**). To test the model, we asked whether ectopic BRD4-NUT expression in non-NC cells sensitizes BRD4L to degradation when treated with BD2i. For this experiment, we used our inducible isogenic 293TRex and U2OSTRex cell lines. Expression of BRD4-NUT in these cells re-localizes BRD4L to dense, BRD4-NUT-containing nuclear condensates corresponding to megadomains, based on immunofluorescence (**Supplementary Fig. S5**). Indeed, BRD4-NUT induced in 293TRex and U2OSTRex cells markedly reduces BRD4L compared with uninduced cells treated with BD2i (DC-9476 and ABBV-744) (**Fig. 4N-O**). This enhanced reduction of BRD4L was not observed with BD1i or BD1-BD2-BETi treatment. These findings support our model that BRD4 must be at a high enough concentration for efficient degradation by SPOP, and offer a possible reason for the lack of sensitivity of non-NC cells to BD2i. Moreover, the model also explains why BRD4-NUT lacking the SPOP degron may protect BRD4L from degradation (**Fig. 4I**). We propose that degron-deficient BRD4-NUT dilutes the concentration of SPOP substrate within megadomains exposed to BD2i, thereby impairing cooperative SPOP assembly and protecting neighboring BRD4L from degradation. Although this model is supported by our findings, further studies will be required to directly establish the proposed role of substrate density in promoting SPOP oligomerization and efficient degradation.

Taken together, the overall findings support our conclusion that SPOP-mediated degradation of BRD4L and BRD4-NUT is required for BD2i to inhibit NC growth, and challenges the prevailing concept that BETi generically acts solely through displacement of these proteins from chromatin^18,20,73^. It is not surprising that this mechanism has remained unknown until now. Because BRD4L is not degraded to the same degree in non-NC cells by BD2i, and because it is not degraded at all by BD1-BD2-BETi, it has not previously been appreciated that BD2i leads to its degradation^27,29,30^. In fact, BET PROTACs were designed^74^ based on the idea that BET inhibitors on their own only displace BRD4, not degrade it^73^. The discovery that BD2i promotes not only BRD4-NUT, but also BRD4L degradation in NC helps explain the unique activity of this class in NC, while also providing mechanistic rationale for clinical development in this cancer and possibly others that harbor high concentrations of chromatin-bound BRD4L.

### BD2-selective inhibition depletes megadomains of BRD4L, BRD3-, and BRD4-NUT

Having found that BRD4L and BRD4-NUT remain chromatin-associated during BD2 inhibition despite undergoing rapid degradation, we next examined the effects of DC-9476 on their genome-wide chromatin occupancy, as well as that of BRD3-NUT. Degradation of BRD4L and BRD4-NUT is expected to deplete their chromatin-bound pools. In contrast, because BRD3-NUT is not degraded in the presence of low-dose DC-9476 (**Fig. 2D**), changes in its chromatin distribution are expected to reflect altered site-specific chromatin occupancy rather than protein loss.

Given that BRD-NUT oncoproteins bind chromatin to form megadomains, we compared the effects of BD2 inhibition by DC-9476 on these massive domains in NC cells with those on enhancers in non-NC cells. For epigenetic analysis, we performed Cleavage Under Targets and Release Using Nuclease (CUT&RUN) on cells treated for 18h with DC-9476 (300nM) using antibodies to wild type BRD4L, NUT, and H3K27ac to identify megadomains, large domains (i.e., super-enhancers), and regular domains (i.e., regular enhancers). Domains in this study are delineated by H3K27ac occupancy (overall size and signal). Large (i.e., megadomains and super-enhancers) or small H3K27ac domains (i.e. regular) are defined, respectively, by being larger or smaller than the point at which a rank-ordered list of region size rapidly increase (i.e., the inflection point of the curve) for H3K27ac modifications^52^. After region size is evaluated, regions with very high signal are retrieved even if they don’t meet the size cut off. Megadomains are defined as H3K27ac domains above the cut-offs for size and signal and also co-enriched with NUT in NC (ranges: 10-15 = 15.4Kb (from signal) to 1.5Mb; 10326 = 14.2Kb (from signal) to 1.1Mb). While megadomains are a subset of super-enhancers, for this CUT&RUN analysis we restrict the term “super-enhancers” for large H3K27ac domains identified without a NUT peak in the given cell type.

Our first question was whether BRD4L is associated with BRD4- or BRD3-NUT megadomains in NC cells, as predicted by proteomic analyses^9^, and also by the fact that BRD4Ls bromodomains are identical to those of BRD4-NUT, and highly homologous to those of BRD3-NUT. Thus, BRD4L would be expected to bind the same acetyl-histone residues and associated genomic regions.

Indeed, while BRD4L is present in domains outside of megadomains, it is present within nearly all megadomains identified in both BRD4- (331/333) and BRD3-NUT (338/340) cell types (**Fig. 5A,E**), even though regions were not selected on the basis of binding to the BRD4L antibody. The re-localization of BRD4L to BRD4-NUT megadomains can be visualized by immunofluorescence in U20STRex and 293TRex cells induced to express BRD4-NUT (**Supplementary Fig. S5**). Whereas BRD4L is diffusely distributed on chromatin at baseline, when BRD4-NUT is expressed, it redistributes to be mostly present within BRD4-NUT foci/megadomains. This also supports our model, which requires that megadomains formed by BRD4-NUT increase the local concentration of BRD4L and BRD4-NUT to increase the efficiency of SPOP-mediated degradation (**Fig. 4M**).

BRD4-NUT+ 10-15 and BRD3-NUT+ 10326 NC cells treated with DC-9476 demonstrated depletion of BRD4L, BRD4-NUT, and BRD3-NUT from megadomains, regular enhancers (**Fig. 5B-C, F-G**), and transcriptional start sites (TSS, not included in figure). These changes correspond with an increase in total number of domains (35% in 10-15s, and 20% in 10326s), most of which are small regions consisting of regular enhancers and TSSs [compare number of regions below the dashed red line pre- (red curve) and post-treatment (blue curve), **Fig. 5D,H**]. The increased numbers of small domains might represent treatment-induced fragmentation of megadomains into smaller domains, and/or an increase in number of new small domains arising from the displacement of BRD-NUT and associated p300.

Compared with NC cells, the super-enhancers in HCC95 cells were smaller than super-enhances or megadomains in NC cells (compare red curves and dashed red lines in **Fig. 5D,5H** with **5L**). As expected, BRD4L was detected in H3K27ac-enriched super-enhancers in HCC95 cells (**Fig. 5I-J**). Whereas H3K27ac signal increased in H3K27ac super-enhancers upon treatment with DC-9476, the BRD4L signal detected in these domains and regular domains was mostly unchanged upon treatment (**Fig. 5K**). These changes in signal did not correlate with a large change in total number of domains (only 3.5% fewer, **Fig. 5L**), which, together with minimal changes in BRD4L enrichment, explains the lack of transcriptional and phenotypic changes observed in non-NC cells.

Taken together, these findings reveal that inhibition of BD2 alone is sufficient to deplete BRD3- or BRD4-NUT and BRD4L from chromatin in NC, an unprecedented finding.

### Chromatin domain size changes induced by BD2 inhibition are associated with concordant transcriptional changes

Enhancers regulate transcription, and when acetylated, are detectable by an increase in the p300/CBP specific mark, H3K27ac, with a frequent concomitant increase in transcription of associated genes. We used overlap or proximity to identify genes potentially affected by DC-9476. In this case, the size of the domains is somewhat confounding since each region, especially megadomains, may overlap multiple genes and enhancers. We correlated H3K27ac domain size with associated gene transcription across all time points measured (18, 24, 36, and 48hrs of treatment). We required that a given gene was consistently up- or down-regulated at all time points. As predicted, H3K27ac domains that decreased in size or were lost mostly correlated with decreased transcription of nearby genes in both BRD4- and BRD3-NUT NC cells (**Fig. 5M-N**). As expected for a non-NC, DC-9476-insensitive control, HCC95 cells showed minimal epigenetic and transcriptional responses to treatment.

### BD2 inhibition-induced megadomain depletion is associated with downregulation of MYC in NC

We next investigated the epigenetic and transcriptional effects of BD2i on key BRD-NUT target proto-oncogenes. As predicted, megadomains were identified at previously described NC-associated lineage-specific or proto-oncogene loci, including *MYC, SOX2, TP63*, in both BRD4- and BRD3-NUT NC cell lines (**Fig. 5O**). Upon DC-9476 treatment, megadomains at all of these loci became depleted of H3K27ac, BRD3/4-NUT, and BRD4L (**Fig. 5O**). Despite these changes, only *MYC* or *MYC*-associated lncRNA expression decreased significantly over the 48h time course, correlating with the observed decrease in MYC protein in these cells (**Fig. 1E, 2D**). Importantly, BRD4/BRD4-NUT expression was unchanged by DC-9476 treatment (**Fig. 4B**).

### BD2 inhibition is more effective than BD1-BD2-BETi at blocking NC growth and prolonging survival, in vivo

Our interest in BD2i stems from their superior safety profile compared with first generation, BD1-BD2-BETi^29,30,75^. The first-in-human study investigating ABBV-744 revealed a low frequency of dose-limiting toxicities (13%)^75^, and lacked the thrombocytopenia or gastro-intestinal toxicity observed frequently in patients treated with BD1-BD2-BETi^21–24,76^. Greater tolerability allows continuous dosing needed for NC patients, and higher doses to facilitate better target engagement. Thus, we compared the pre-clinical efficacy of BD2i (ABBV-744 and DC-9476) with that of the BD1-BD2-BETi, ABBV-075.

In our first study, we compared the activity of ABBV-744 with that of ABBV-075 in our 10-15 luciferized, disseminated xenograft model of NC, using a similar approach to that described above (see **Fig. 1D**). After disseminated tumor was established based on bioluminescence (BLI) signal ≥ 2×10^6^ ph/sec/cm²/sr, on day 7, treatment began and continued for only 28 days, after which mice were observed for durability of treatment effect and survival. Strikingly, equivalent activity was seen comparing the maximum tolerated dose (MTD) of ABBV-075 (1mg/kg) with only 1/16^th^ the MTD of ABBV-744 (4.7mg/kg)^29^, with slight prolongation of survival in both cohorts (**Fig. 6A**). However, tumor growth repression was markedly enhanced, and survival was significantly prolonged in animals treated with the MTD of ABBV-744 (75 mg/kg) compared with those treated with ABBV-075.

**Fig. 6.**
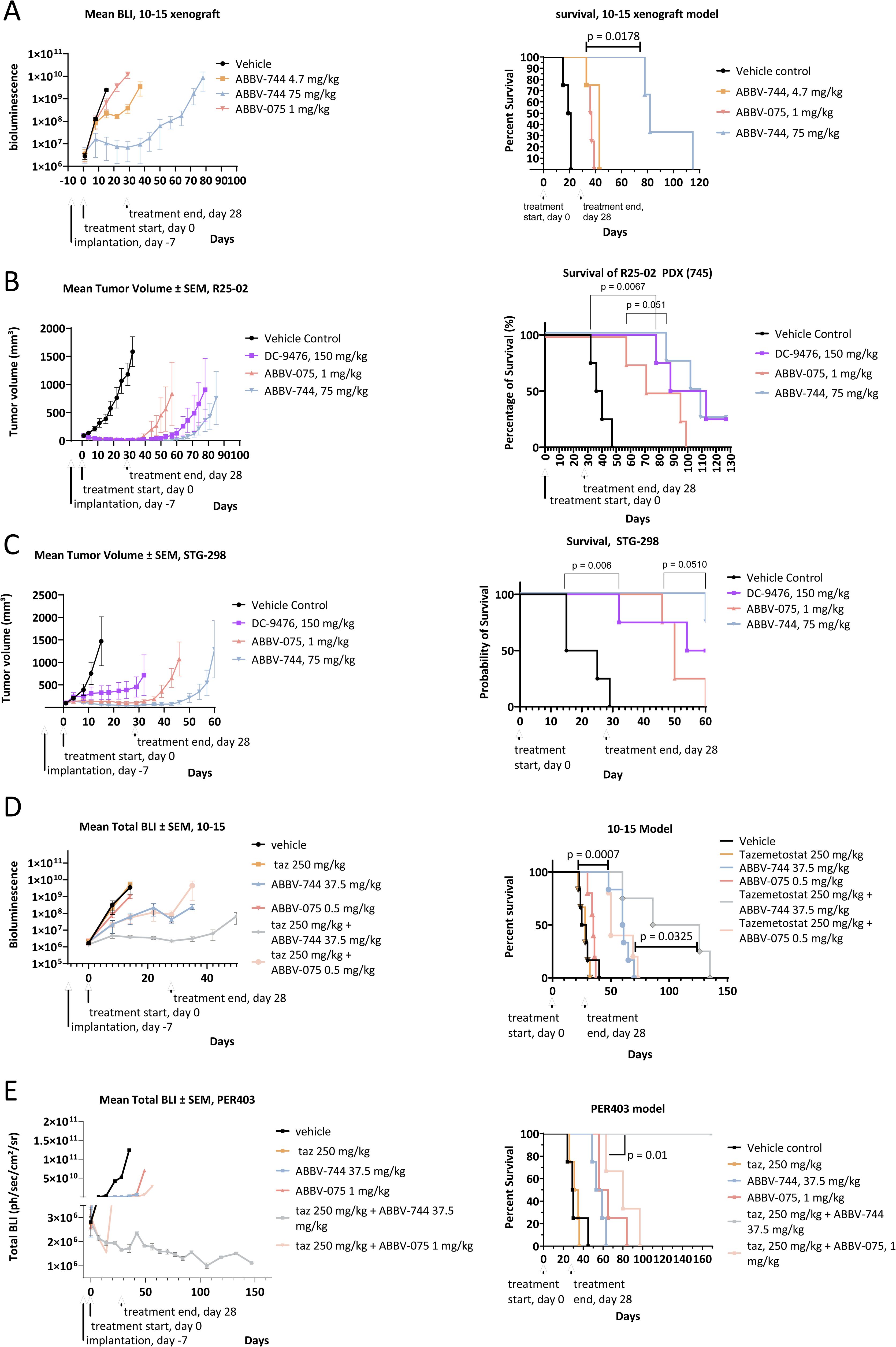
BD2 inhibition is more effective than BD1-BD2-BETi at blocking NC growth and prolonging survival in vivo. **A-E**. (Left) Tumor growth over time measured by total bioluminescence (BLI) or tumor volume. Treatment is initiated on day 0 and is continuously administered until treatment termination on day 28. (Right) Kaplan-Meier survival plots corresponding to left. In **D-E**, vehicle, ABBV-744, tazemetostat, and ABBV-744 plus tazemetostat cohorts were previously reported in Huang et al.^18^ and are included here as re-plotted published data for comparison with unpublished ABBV-075 ± tazemetostat cohorts from the same in vivo studies. **F**. Mechanistic model of BD2i-induced degradation of BRD4L and BRD4-NUT by SPOP-CUL3.

To determine whether similar differences are seen comparing the MTD of ABBV-744 and DC-9476 with ABBV-075, we utilized a new, BRD4-NUT+ patient-derived xenograft model (PDX) of NC, R25-02, implanted in the flanks of NSG mice. This PDX was obtained from an eighteen-year-old female who died from a BRD4-NUT+ NC of the thyroid. After tumor was established based on caliper measured volume of ≥ 90mm^3^, on day 7, treatment began and was continued for 28 days then terminated. As with cell line derived xenografts, vehicle-treated tumors grew rapidly and all mice died within 47 days of the treatment start date. BETi treatment with all three compounds significantly prolonged survival compared with vehicle treatment (p = 0.0001) (**Fig. 6B)**, and all compounds were tolerated well based on lack of weight loss (**Supplementary Fig. S6A**). Importantly, tumor regression was observed in all BETi-treated mice. However, similar to the previous experiment, there was greater tumor growth suppression and prolonged survival in mice treated with the BD2i, ABBV-744 and DC-9476, than those treated with the BD1-BD2-BETi, ABBV-075.

To determine whether BD2i, particularly the BRD4-BD2-selective DC-9476, is effective in treating BRD3-NUT+ NC, we pre-clinically evaluated these compounds and ABBV-075 in a PDX flank implantation model, STG-298. This PDX was obtained from a surgical resection of a nasal NC in a 29-year-old male. As in the previous PDX, when tumor was established based on caliper measured volume of ≥ 100mm^3^, on day 7, treatment began and was continued for 28 days then terminated. Similar to R25-02, vehicle-treated tumors grew rapidly and all mice died within 29 days of the treatment start date. BETi treatment with all three compounds significantly prolonged survival compared with vehicle treatment (p < 0.0001) (**Fig. 6C**). As with R25-02, all compounds were tolerated well based on lack of weight loss (**Supplementary Fig. S6B**). In contrast to the BRD4-NUT+ PDX, slight tumor regression and durable growth inhibition was observed only in BD2i-(ABBV-744) treated mice, whereas deaths due to tumor progression began in both ABBV-075 and DC-9476-treated mice within 18 days following treatment termination. Notably, DC-9476 was less active than either ABBV-744 or ABBV-075, consistent with its 9-fold lower affinity for BRD3 BD2 relative to BRD4 BD2 (**Table 1**). The findings support the concept that BD2i has superior therapeutic efficacy in treatment of NC than BD1-BD2-BETi, but importantly suggest that non-BRD4-NUT+ tumors may be less responsive to BRD4-BD2-selective inhibition by DC-9476. The favorable safety profile of BD2i is predicted to allow for greater tolerance to combinations with other targeting agents using effective doses. In fact, we previously showed that ABBV-744 combined with tazemetostat can lead to durable complete responses in our disseminated xenograft models of NC^18^. However, the efficacy of this combination was not compared with that of a BD1-BD2-BETi and tazemetostat. Thus, to directly compare the activity of these two combinations in the same in vivo experiment, we analyzed previously unpublished ABBV-075 ± tazemetostat cohorts alongside vehicle, ABBV-744, tazemetostat, and ABBV-744 + tazemetostat cohorts that were reported previously^18^. In two of three models (10-15 and PER-403), there was markedly improved tumor growth repression in ABBV-744 + tazemetostat-treated animals than in those treated with ABBV-075 + tazemetostat (**Fig. 6D-E**). Moreover, in both of these models there was significantly improved survival in ABBV-744 + tazemetostat cohorts compared with those treated with ABBV-075 + tazemetostat [p = 0.01 (PER-403) and 0.0325 (10-15)]. Importantly, long term complete responses were only seen in BD2i + tazemetostat-treated mice.

Collectively, these in vivo studies with structurally diverse BD2-selective inhibitors demonstrate that improved tolerability permits higher exposures than are achievable with BD1-BD2-BET inhibitors, thereby enabling more complete target inhibition and potentially curative combination regimens in NC.

## Discussion

The data above provide a new mechanistic understanding of how BD2i inhibits NC growth. The prior model viewed SPOP as a passive homeostatic regulator of BRD4 abundance, even in the presence of BETi^62,68,69^. Our data suggest instead that BD2 inhibition actively converts BRD4L and BRD4-NUT into substrates for SPOP-mediated proteasomal degradation (illustrated in **Fig. 4M**). Our data further demonstrate that SPOP-CUL3–mediated degradation is an essential component of BD2 inhibitor activity, rather than a dispensable downstream consequence of chromatin displacement. This was demonstrated by SPOP knockout, which nearly completely prevented BD2i-induced phenotypic changes in NC cells, including MYC downregulation, differentiation, and cell-cycle arrest (**Fig. 4A-F**). The observation that BD2i induces BRD4-NUT loss in preclinical NC models further supports the therapeutic relevance of this degradation mechanism (**Fig. 2F**).

Our data also support a mechanistic model **(Fig. 4M**) that may explain the much greater sensitivity of NC compared with non-NC cells to BD2i. in this model, BRD4-NUT creates a favorable environment for efficient SPOP-mediated degradation of itself and BRD4L in the presence of BD2i, as follows. BRD4-NUT forms hyperacetylated chromatin megadomains endogenously and when ectopically expressed^16,77^. The tandem bromodomains of BRD4L and BRD4-NUT engage in multivalent binding to acetylated histones, with affinity increasing with the number and density of acetyl marks, such as di-acetylated histone H3 K9/14 or histone H4 K5/12^26^ ^77–79^. These densely clustered, multivalent histone acetyl marks bind BRD4L and BRD4-NUT bromodomains with high affinity^7^, thereby accumulating large numbers of closely spaced BRD4L and BRD4-NUT molecules, as seen by immunofluorescence (**Fig. 4H, Supplementary Fig. S5**) and CUT&RUN (**Fig. 5**). It has been shown that multivalent substrates promote higher-order SPOP assemblies and enhance SPOP-mediated ubiquitination, while increasing concentrations of SPOP and substrate favor formation of these higher-order assemblies^71,72^. While BRD4L and BRD4-NUT do not possess multivalent SPOP-degrons, we hypothesize that the high local concentration of BRD4L and BRD4-NUT within megadomains creates a substrate environment that is effectively multivalent. Thus, BD2i-induced degron exposure within megadomains generates a high-density array of SPOP-binding sites that facilitate higher-order SPOP assembly and efficient ubiquitination of BRD4 and BRD4-NUT. Because non-NC cells lack the BRD4-NUT-driven megadomains characteristic of NC, they may not produce this same high-density substrate environment upon BD2i treatment. This relative resistance to SPOP-mediated degradation can be overcome by ectopic BRD4-NUT expression (**Fig. 4N-O**).

The unique sensitivity of NC to BD2 inhibition is particularly favorable for clinical development because antitumor activity can be achieved without BD1 inhibition, whose chromatin-binding function is required for transcription of essential housekeeping genes in healthy cells^25–27^. The greater tolerability of BD2i allows for higher dosing, greater target engagement, and was demonstrated here to have superior pre-clinical activity compared with BD1-BD2-BETi in BRD4-NUT+ models and in our BRD3-NUT+ model (**Fig. 6**). Moreover, the favorable toxicity profile of BD2i may enable combinations with other targeted agents that are more effective and better tolerated than combinations with BD1-BD2-BETi, as evidenced by several of our models (**Fig. 6D-E)**. This is important because combination approaches will likely be needed to fully address NC biology.

If validated, our mechanistic model explains why NC is uniquely sensitive to BD2i and may help identify non-NC cancers that are also sensitive to BD2i. Other BRD4-dependent tumors in which BRD4 is unusually concentrated on chromatin, or in which SPOP activity is enhanced, may also be BD2i-sensitive. One example is the 4-10% of endometrial cancers with gain-of-function SPOP mutations^68,80^. It has been shown that endometrial adenocarcinomas with gain-of-function SPOP mutations have reduced baseline levels of BRD4L and are consequently more sensitive to BETi^68^. However, this study was reported in 2017, before the development of BD2i, and it will therefore be important to determine whether these tumors are also more sensitive to BD2i.

Androgen-receptor (AR)-dependent prostate cancer has previously been shown to be uniquely BD2i-sensitive^29^. In that study, sensitivity was attributed to eviction of BRD4 specifically from AR-containing super-enhancers. The effect of BD2i on BRD4 protein levels was not evaluated, nor was a distinction made between SPOP-mutated and wild-type tumors. Separately, loss-of-function mutations of SPOP (6-15% of all prostate cancers^81^) were shown to increase baseline levels of BRD4 and confer resistance to BETi^62,69^. Our findings warrant testing whether BD2i resistance in AR-dependent prostate cancer is conferred by impaired SPOP-dependent degradation of BRD4 caused by loss-of-function SPOP mutations. Conversely, sensitivity of these tumors to BD2i may result from partial inhibition of BRD4 function and/or BD2i-induced BRD4 degradation.

There are several limitations of the presented work. First, additional studies are required to validate our model, which at this stage remains theoretical until it is demonstrated that the high local concentration of substrate BRD4L and BRD4-NUT in NC enhances SPOP assembly and activity following BD2 inhibition. In particular, BD2i-induced higher-order SPOP assembly remains to be demonstrated directly. Moreover, the predictive value of high-density chromatin-associated BRD4 or SPOP gain-of-function mutations for BD2i sensitivity in non-NC cancers needs to be tested. One potential challenge to our model is our finding that 24h treatment with BD1-BD2-BETi induces loss of BRD4-NUT in NC cells (**Supplementary Fig. S3**). However, this occurred with delayed kinetics, not within 2h (**Fig. 4K**, left panel), and without corresponding loss of BRD4L, suggesting a mechanism distinct from the rapid, dual degradation produced by BD2-selective inhibition.

Finally, this work introduces a new BRD4-selective BD2i, DC-9476, originally developed for inflammatory disease, as a potent therapeutic candidate for NC. DC-9476 has recently been granted orphan drug designation by the FDA for treatment of NUT carcinoma (DRU-2025-10835).

## ACKNOWLEDGEMENTS

DC-9476 was generously provided by DeepCure. DeepCure reviewed this manuscript for scientific accuracy, and provided data they generated in creating DC-9476. ABBV-075 and ABBV-744 were graciously provided by ABBVIE, which also reviewed this manuscript for scientific accuracy. In addition, Ipsen kindly provided tazemetostat for this and the previously reported study^18^, and has also reviewed this manuscript for scientific accuracy.

## Author contributions

K.A.B. performed experiments, analyzed data, supervised experiments, provided intellectual input and prepared figures; H.N. performed experiments, analyzed data, and prepared figures; K.P.E. provided intellectual input and developed methods; J.H. performed experiments and analyzed data; P.C.G. supervised animal studies and helped interpret data; B.A.L. performed animal studies and analyzed the resulting data; B.K.E. performed animal studies and analyzed the resulting data; S.T.G. provided essential reagents; K.S. provided bioinformatic methods; R.T.D. performed experiments and provided methods; J.L. provided intellectual input; G.I.S. provided intellectual input; S.J.F. led the medicinal chemistry development of DC-9476, including compound design, synthesis, and optimization, and contributed to analysis of its biochemical selectivity and pharmacologic properties; J.H.G. contributed to project administration, resources, and supervision. D.R. performed experiments and analyzed data; K.R.S. contributed to resources, funding acquisition, project administration, and review and editing of the manuscript; L.R. contributed to conceptualization, supervision, project administration, resources, and review and editing of the manuscript; M.E.L. performed bioinformatic analysis of RNAseq and CUT&RUN data; C.A.F. conceived and supervised the study, obtained funding, interpreted the data, performed experiments, analyzed data, prepared figures, and wrote the manuscript. All authors reviewed and edited the manuscript.

