## Supplementary Figures S1-S6 and Tables S1-S5 for "BET BD2 inhibition facilitates SPOP-mediated degradation of chromatin-associated BRD4/BRD4-NUT, a therapeutic vulnerability in NUT carcinoma"

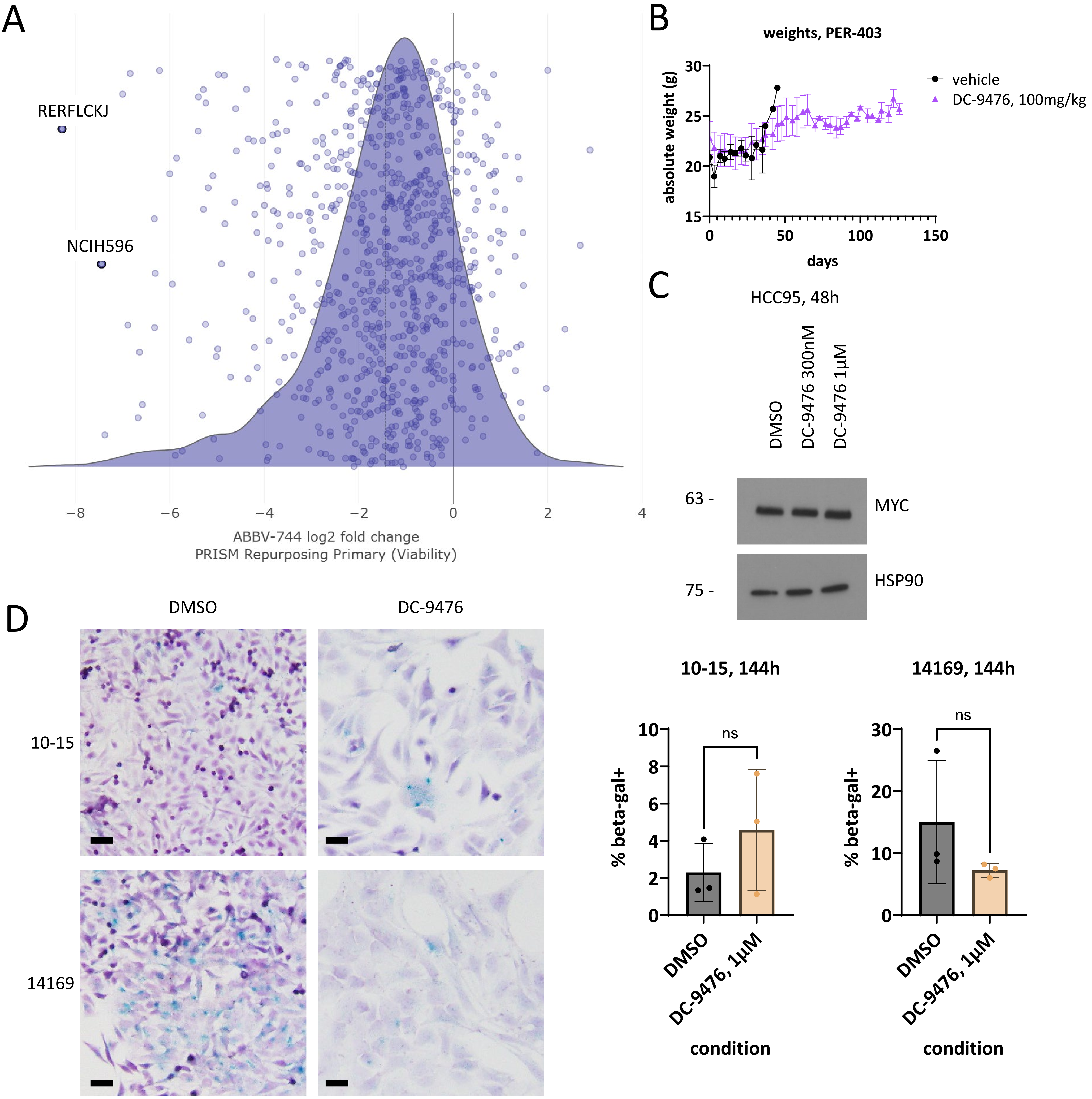


**Supplementary Fig. S1.** BD2-selective inhibition blocks growth of a subset of diverse cancer cell types, does not affect MYC expression in non-NC cells, is well tolerated in mice, and has a variable effect on senescence. **A**. DepMap drug sensitivity analysis as indicated. **B.** Weights over time for treatments indicated. N = 4 mice per arm. **C**. Immunoblot of non-NC HCC95 cells treated with DC-9476 for 48h as indicated. **D.** (Left) Images of indicated cell lines stained for beta-galactosidase (blue). Counterstain, hematoxylin. Scale bars, 50μm. (Right) Comparison of beta-galactosidase staining based on cell counts of three biologic replicates. Comparison was made by two-tailed paired Student's t-test, with each independent experiment constituting a matched pair. Individual biological replicate values are shown together with the mean ± SD.

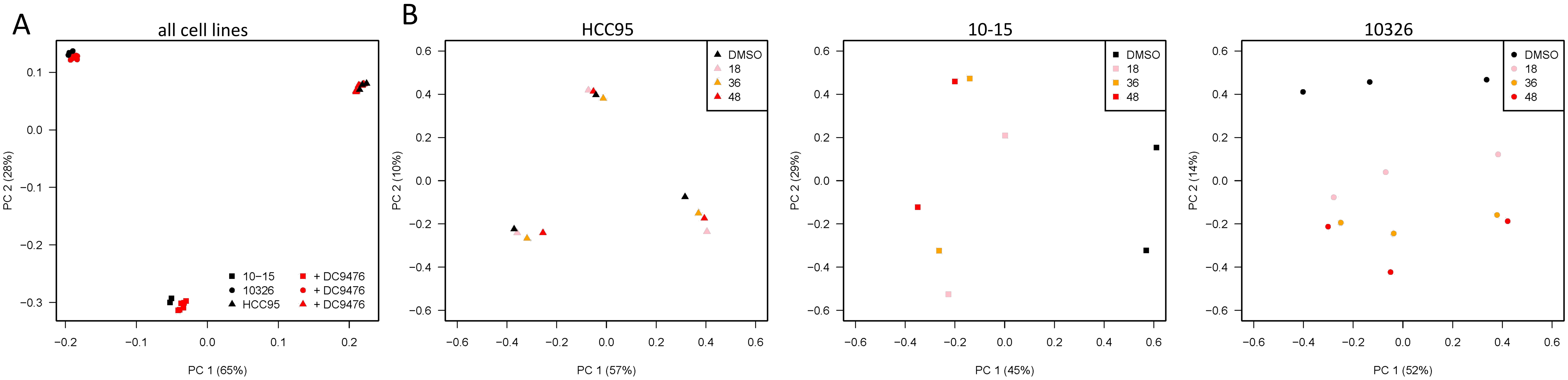


**Supplementary Fig. S2.** Principal component analysis of RNAseq on cells treated as indicated based on top 500 genes by variance. **A.** All cell lines. **B.** Individual cell lines.

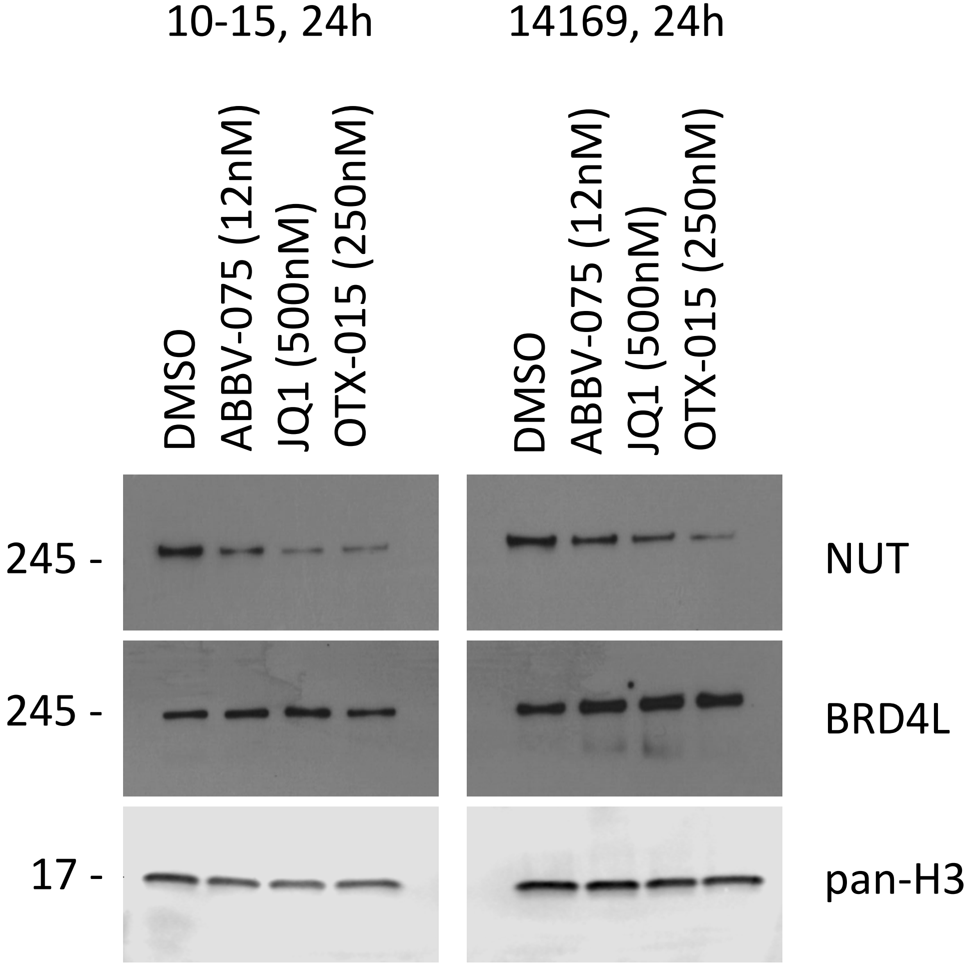


**Supplementary Fig. S3.** BD1-BD2-BET inhibition depletes BRD4-NUT but not BRD4L in NC. Immunoblot of NC cells treated with diverse BD1-BD2-BETi as indicated.

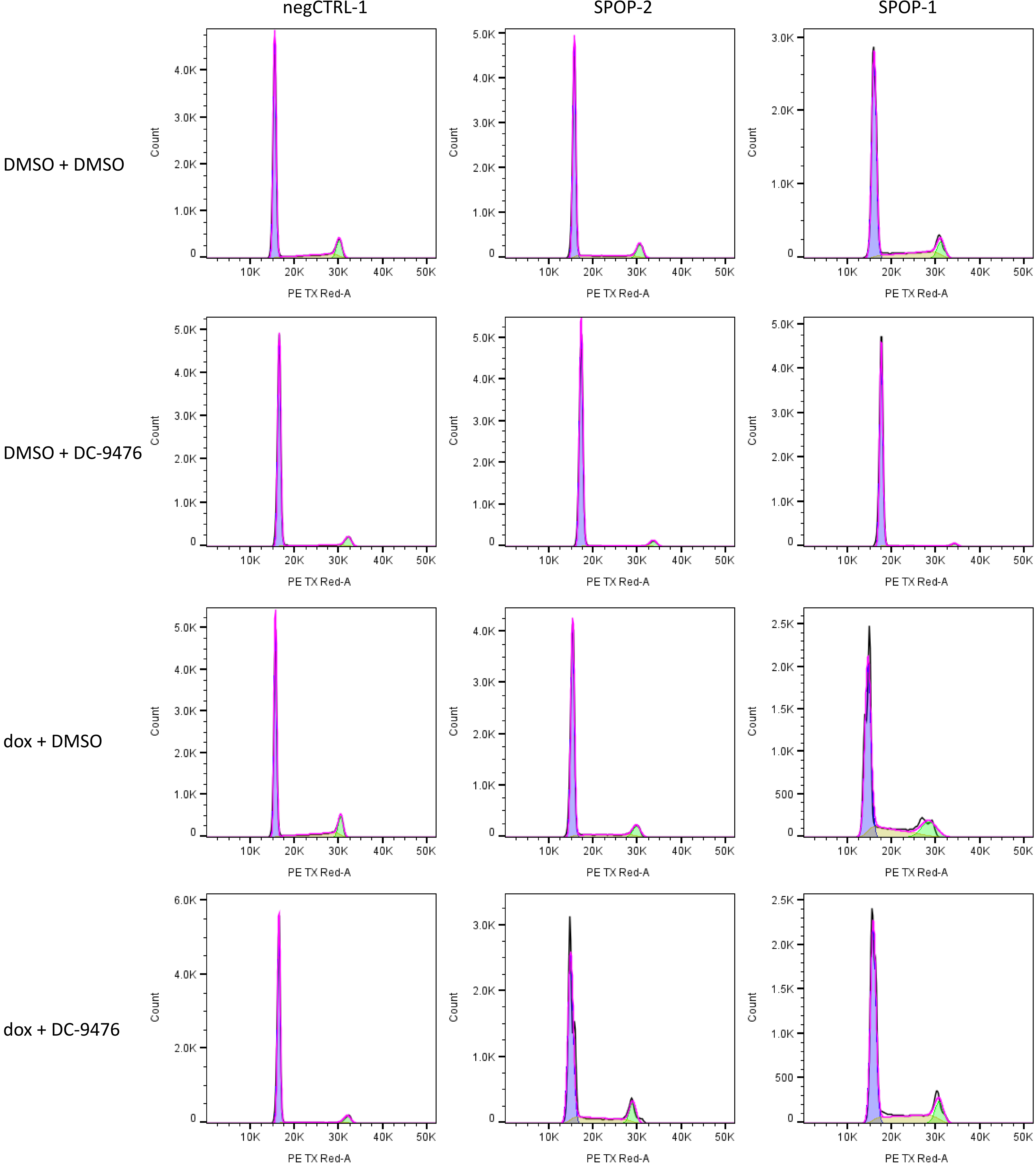


**Supplementary Fig. S4.** Histograms of representative single replicates corresponding to cell cycle analysis performed in triplicate in **Fig. 4E.**

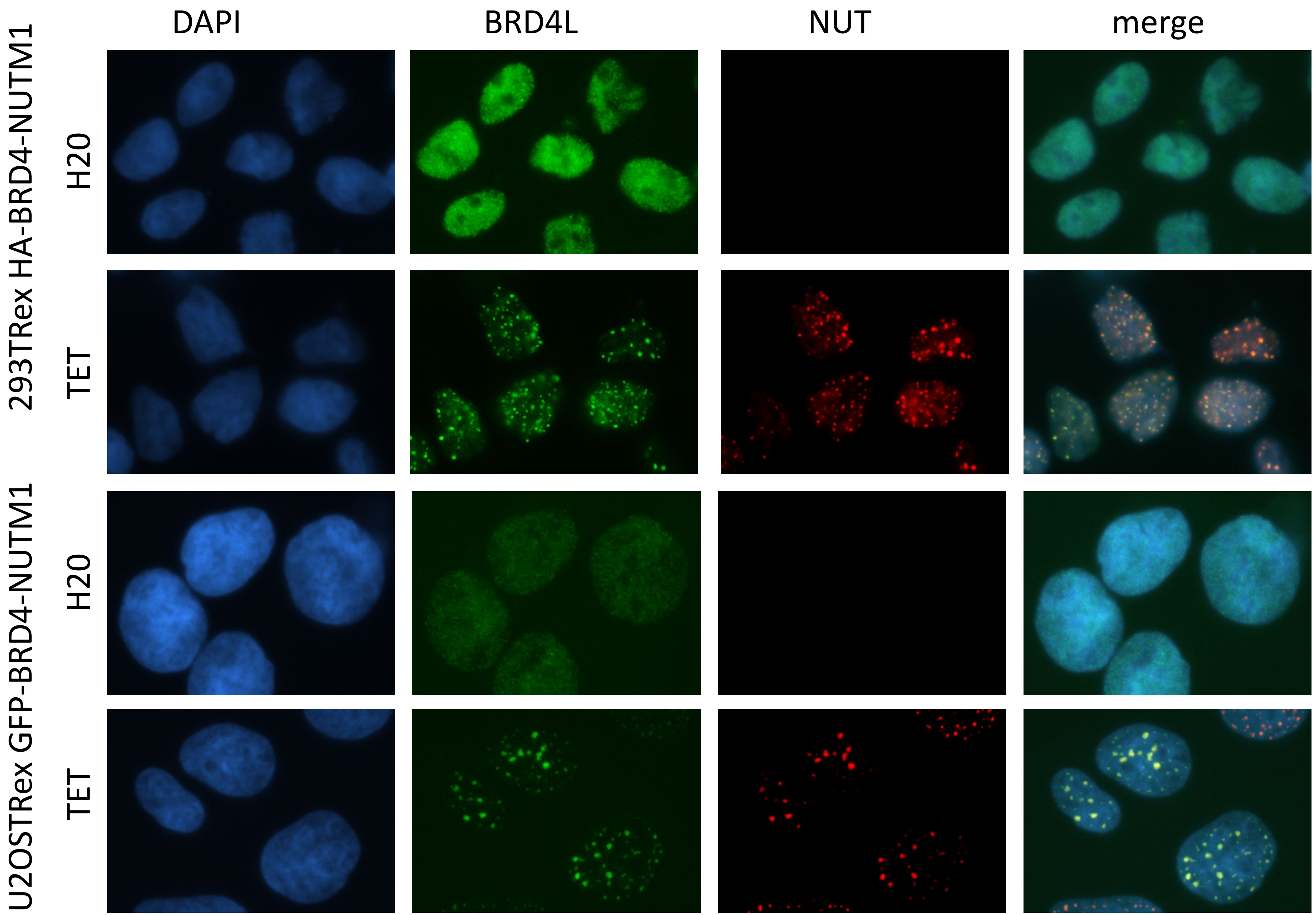


**Supplementary Fig. S5.** Expression of BRD4-NUT in non-NC cells re-localizes BRD4L to nuclear condensates corresponding to megadomains. Immunofluorescence microscopy using antibodies as indicated.

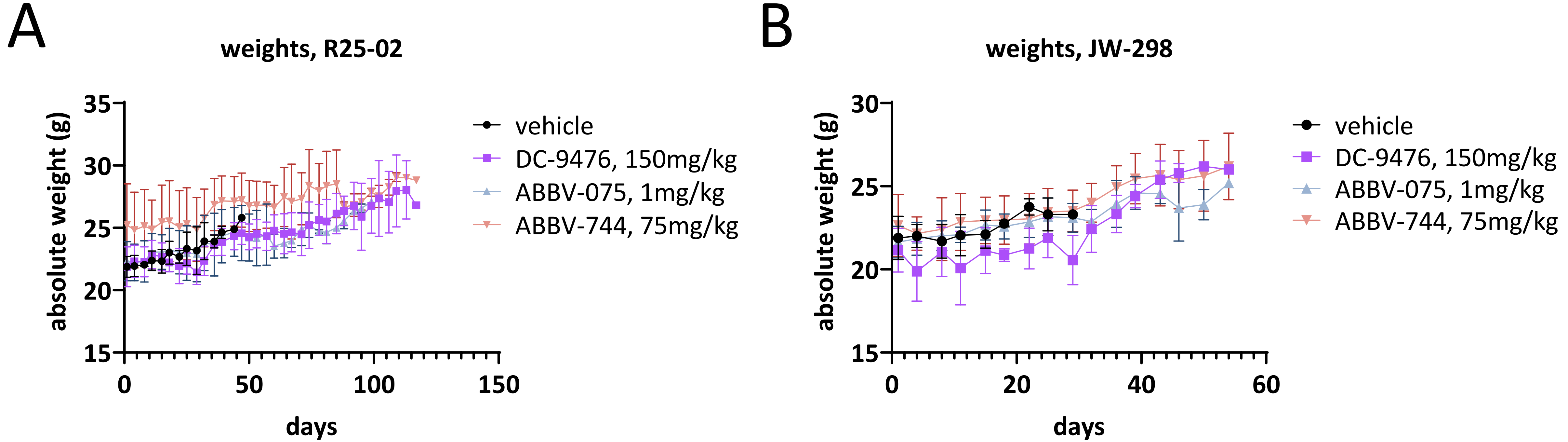


**Supplementary Fig. S6.** Treatment with BD2i or BD1-BD2-BETi does not lead to weight loss exceeding 10% of body weight in NSG mice. **A-B.** Weights over time for treatments and models indicated. N = 4 mice per arm.

| **Supplementary Table S1.** antibodies used for immunohistochemistry. | | | | | | |
|---|---|---|---|---|---|---|
| protein recognized | dilution | catalogue no. | clone | company | detection | antigen retrieval |
| NUT | 1:100 | 3625 | C52B1 | CST | Leica Bond – Leica Refine Detection Kit (DS9800) | |

| **Supplementary Table S2.** Primary antibodies used for immunoblotting. | | | | |
|---|---|---|---|---|
| protein recognized | dilution | catalogue no. | clone | company |
| KRT7 (CK7) | 1:1,000 | 15539-1-AP | rabbit polyclonal | ProteinTech |
| involucrin | 1:1000 | I9018 | SY5 | Millipore Sigma |
| histone H3 | 1:4,000 | ab1791 | rabbit polyclonal | CST |
| NUT | 1:1,000 | 3625 | C52B1 | CST |
| PARP | 1:1,000 | 9542 | rabbit polyclonal | CST |
| BRD4L | 1:5,000 | A301-985A | rabbit polyclonal | Bethyl Laboratories |
| HSP90 | 1:5,000 | 4877 | C45G5 | CST |
| c-myc | 1:1,000 | 5605 | D84C12 | CST |
| SPOP | 1:1,000 | 68216-1-Ig | 2B9F2 | ProteinTech |

| **Supplementary Table S3.** Secondary antibodies used for immunoblotting. | | | |
|---|---|---|---|
| Species recognized | dilution | catalogue no. | company |
| IRDye® 800CW Goat anti-Rabbit IgG Secondary Antibody | 1:10,000 | 926-32211 | LICOR Bio |
| Anti-mouse IgG, HRP linked Antibody | 1:1,000 | 7076S | Cell Signaling Technology |
| Anti-rabbit IgG, HRP linked Antibody | 1:1,000 | 7074S | Cell Signaling Technology |

| **Supplementary Table S4.** Primary antibodies used for immunofluorescence. | | | | |
|---|---|---|---|---|
| protein recognized | dilution | catalogue no. | clone | company |
| H3K27ac | 1:1,000 | 39685 | MABI 0309 (mouse) | Active Motif |
| NUT | 1:1,000 | 3625 | C52B1 | CST |
| BRD4L | 1:2,000 | 63759 | E4X7E | CST |

| **Supplementary Table S5.** Secondary antibodies used for immunofluorescence microscopy. | | | |
|---|---|---|---|
| Species recognized/Wavelength | dilution | catalogue no. | company |
| Anti-mouse Alexa Fluor 488 | 1:2,000 | A-11001 | Invitrogen |
| Anti-mouse Alexa Fluor 594 | 1:1,000 | A-11005 | Invitrogen |
